# Optogenetic low-frequency stimulation of dentate granule cells prevents seizure generation in experimental epilepsy

**DOI:** 10.1101/2020.01.09.900084

**Authors:** E Paschen, C Elgueta, K Heining, DM Vieira, C Orcinha, U Häussler, M Bartos, U Egert, P Janz, CA Haas

**Author notes:** These authors contributed equally to this work (P Janz, CA Haas). Address of corresponding author: Prof. Dr. Carola A. Haas, Medical Center, University of Freiburg, Dept. of Neurosurgery, Exp. Epilepsy Research, Breisacherstr. 64, 79106 Freiburg, Germany, **Email:**.

## Abstract

Mesial temporal lobe epilepsy (MTLE) is the most common form of focal epilepsy in adults and is typically associated with hippocampal sclerosis and drug-resistant seizures. As an alternative to curative epilepsy surgery, brain stimulation evolves as a promising approach for seizure-interference. However, particularly in MTLE with severe hippocampal sclerosis, current stimulation protocols are often not effective. Here, we show that optogenetic low-frequency stimulation (oLFS) of entorhinal afferents exhibits unprecedented anti-ictogenic effects in chronically epileptic mice. Photostimulation at 1 Hz resulted in an almost complete suppression of focal seizures, independent of the degree of hippocampal sclerosis. Furthermore, by performing oLFS for 30 min before a pro-convulsive stimulus, seizure generalization was successfully prevented. Finally, acute slice experiments revealed a decreased excitability upon oLFS, which may partially explain the observed anti-epileptic effects. Taken together, our results suggest that oLFS of entorhinal afferents constitutes a promising approach for seizure control in MTLE.

## Introduction

Mesial temporal lobe epilepsy (MTLE) represents the most common form of acquired epilepsy in adults and is thought to arise from pro-epileptic modifications of the mesio-limbic network (e.g., the hippocampus and entorhinal cortex) due to an initial precipitating insult (Engel, 2001), which could be a *status epilepticus* (SE), complex febrile seizures or trauma in early childhood. The most frequent histopathological hallmark of MTLE is hippocampal sclerosis, which is characterized by neuronal cell loss and reactive gliosis and is often associated with granule cell dispersion (GCD) and mossy fiber sprouting (Thom, 2014). MTLE is of particular clinical interest since it is often resistant to medication and surgical resection of the epileptic focus represents the only therapeutic option. However, many patients do not remain seizure-free following curative epilepsy surgery (Mohan et al., 2018; Ryvlin & Kahane, 2005), thus demonstrating the urgent need for new therapeutic avenues.

One alternative approach for treating patients with intractable epilepsy is electrical deep brain stimulation. Typically, high-frequency stimulation at 100-200 Hz is performed in the hippocampus or the anterior thalamic nucleus to interfere with limbic seizures (Li & Cook, 2018). However, in MTLE with severe hippocampal sclerosis, electrical stimulation appears remarkably ineffective most likely due to extensive neuronal cell loss and glial scarring (Velasco et al., 2007). To gain a mechanistic and circuit-based understanding of anti-ictogenic effects, electrical stimulation achieves high temporal and spatial control but lacks the necessary cell- or pathway-specificity. Optogenetic modulation of neuronal activity provides the required cell-specificity and has previously been applied to alleviate seizure acidity in several animal studies (Esther Krook-Magnuson, Armstrong, Oijala, & Soltesz, 2013; Xu et al., 2016; Zhao, Alleva, Ma, Daniel, & Schwartz, 2015).

The emergence of seizures is classically attributed to a shifted excitation/inhibition balance, hence the most common approaches for seizure intervention in the hippocampus so far are either based on inhibition of excitatory neurons or the recruitment of GABAergic interneurons (Kokaia, Andersson, & Ledri, 2013; E. Krook-Magnuson, Szabo, Armstrong, Oijala, & Soltesz, 2014; Ladas, Chiang, Gonzalez-Reyes, Nowak, & Durand, 2015; Ledri, Madsen, Nikitidou, Kirik, & Kokaia, 2014; Lu et al., 2016). The latter, however, resulted in an increased seizure probability during optogenetic stimulation (Lévesque et al., 2019). Other approaches investigated the effects of low-frequency stimulation (LFS) of entorhinal principal cells *in vitro* (Shiri et al., 2017) and in a kindling model *in vivo* (Xu et al., 2016), demonstrating promising anti-ictogenic effects on evoked epileptic discharges. Nonetheless, it remains elusive whether LFS is effective in interfering with spontaneous, recurrent seizures in chronic MTLE. This question is of particular importance considering that pyramidal cells and GABAergic interneurons, putatively the main cellular substrate for effective LFS, are lost in the sclerotic hippocampus (Thom, 2014).

In the present study, we addressed this question in mice that had received kainate (KA) into the right dorsal hippocampus to trigger SE and subsequent epileptogenesis. Importantly, this well-established MTLE mouse model recapitulates the major hallmarks of the human pathology, comprising the emergence of spontaneous recurrent seizures and robust unilateral hippocampal sclerosis (Bouilleret et al., 1999) with a sparing of dentate granule cells (DGCs) and CA2 pyramidal cells (Häussler, Rinas, Kilias, Egert, & Haas, 2016) as well as their entorhinal inputs (Janz et al., 2017a). In order to explore the anti-ictogenic effects of pathway-specific optogenetic low-frequency stimulation (oLFS) *in vivo*, we selectively photostimulated entorhinal afferents, which terminate on DGC dendrites in the lesioned hippocampus, during local field potential (LFP) recordings. We present evidence that oLFS is highly effective in preventing both subclinical and behavioral seizures in experimental MTLE with severe hippocampal sclerosis.

## Results

### Variability of hippocampal sclerosis and seizure burden in chronically epileptic mice

First, we characterized the disease severity in mice injected with three different KA concentrations (10, 15 and 20 mM) by quantifying GCD, the extent of cell loss in CA1 and hilus, and the seizure rate.

Quantitative analysis of GCD in NeuN-stained sections 35 to 40 days after KA injection revealed substantial differences between mice injected with low (10 mM) or high (20 and 15 mM) KA concentrations (Figure 2A). The volume of the dispersed GCL (i.e., the extent of GCD), was comparable between 20 and 15 mM KA but significantly less in the 10 mM KA group (Figure 2B, 10 mM: 0.24 ± 0.08 mm^3^; 15 mM: 1.56 ± 0.16 mm^3^; 20 mM: 1.52 ± 0.11 mm^3^, 10 mM vs 15 mM and vs 20 mM p<0.001; n=4/6/5 animals). Conversely, the loss of CA1 pyramidal cells, determined as the total length of cell body-free CA1, was similar in all groups (Figure 2C, 10 mM: 37.72 ± 8.84 mm; 15 mM: 49.57 ± 3.27 mm; 20 mM: 42.28 ± 6.44 mm; n = 4/6/5 animals).

**Figure 1.**
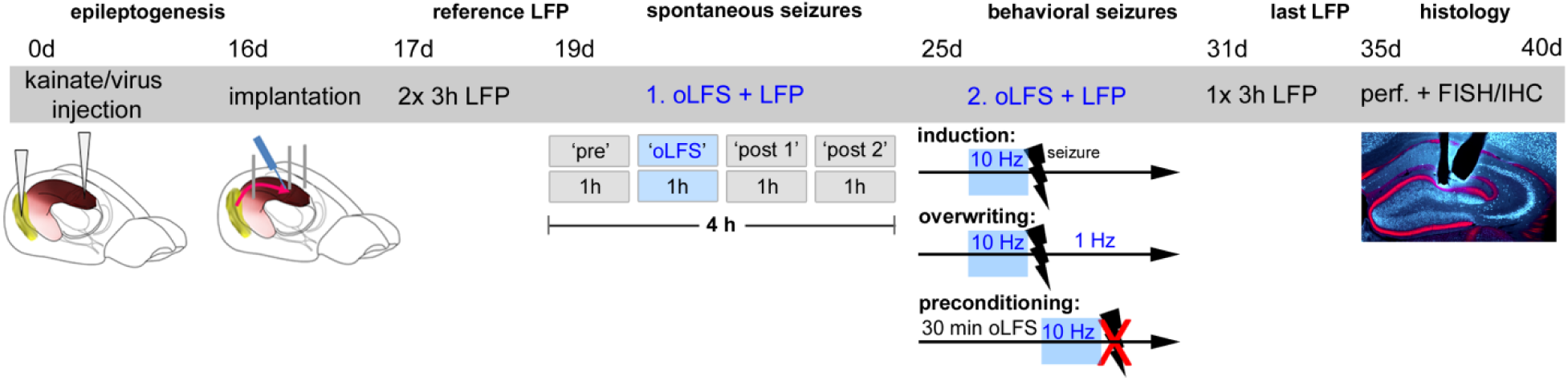
Experimental design for *in vivo* oLFS. Animals received intrahippocampal KA and a ChR2-carrying virus into the entorhinal cortex to trigger epileptogenesis and the expression of ChR2 in entorhinal afferent fibers. At 16 days post injections, electrodes and an optic fiber were implanted. Following recovery from implantations, reference LFPs were recorded on two successive days for three hours each. Next, the effect of oLFS on spontaneously occurring seizures was tested. To this end, recording sessions of four hours per day were performed in all animals. In the first hour (‘pre’), LFPs were recorded to determine spontaneous epileptic activity followed by the application of oLFS pulses for one hour (‘oLFS’). The effect of the optogenetic stimulation was determined by further two hours of recording (‘post 1 and 2’). Three different oLFS frequencies (1, 0.5 or 0.2 Hz) were applied on successive days in each animal (two sessions per animal). In the next block, generalized seizures were induced by optogenetic (10 Hz) stimulation and oLFS (1 and 0.5 Hz) was applied to test potential seizure suppressive effects (overwriting). In addition, a seizure preventing action was tested by applying oLFS prior to the pro-convulsive (10 Hz) stimulation (preconditioning). Finally, animals were perfused after the last LFP recording session and brain sections were processed for immunohistological procedures. FISH, fluorescent *in situ* hybridization; IHC, immunohistochemistry; perf., perfusion

**Figure 2.**
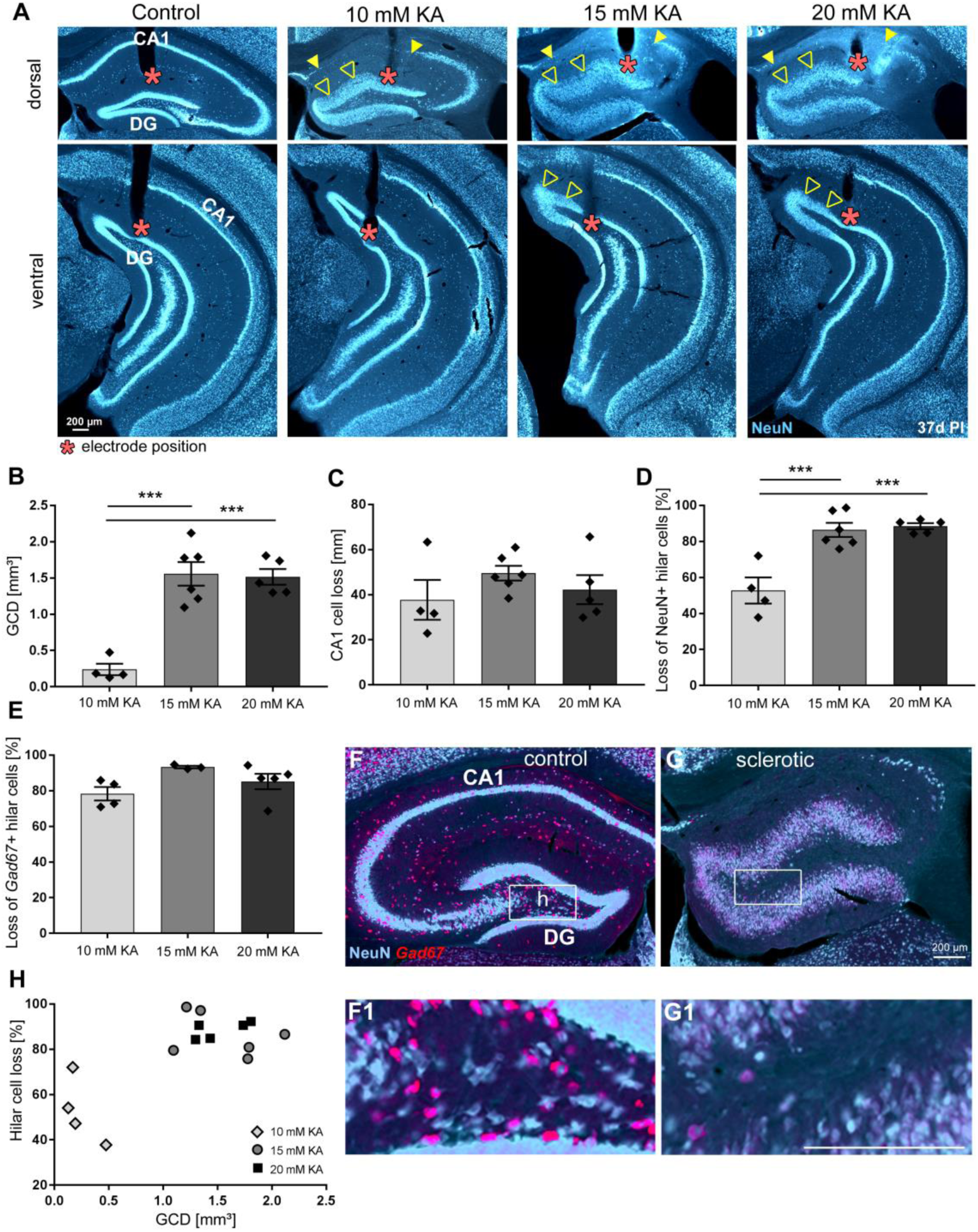
Dependence of hippocampal sclerosis on KA concentration. **(A)** Representative NeuN-labeled sections of dorsal and ventral hippocampal regions treated with different KA concentrations at 37 days post injection (PI). In each section, the electrode position is marked with a red asterisk. Epileptic hippocampi show GCD in the dentate gyrus (open arrowheads) and cell loss in CA1 (region between filled arrowheads). Analysis of hippocampal sclerosis by quantification of **(B)** GCL volume of dispersed regions (i.e., GCD), **(C)** total length of cell loss in CA1, **(D)** % loss of NeuN^+^ hilar cells and **(E)** loss of *Gad67^+^* hilar interneurons in the sclerotic vs non-sclerotic hippocampus (i.e. **(G**, **G1)** ipsilateral vs **(F, F1)** contralateral). One-way ANOVA; Tukey’s multiple comparison test; *p<0.05, **p<0.01 and ***p<0.001. All values are given as mean ± SEM. **(H)** Animals with stronger hilar cell loss (15 and 20 mM KA) show a higher degree of GCD. Scale bars 200 µm.

Comparing the density of hilar neurons in the sclerotic (ipsilateral) and the non-sclerotic (contralateral) hippocampus (Figure 2F, G), we found a significantly smaller loss of NeuN^+^ neurons in the hilus at the KA injection site (idHC) of mice injected with 10 mM KA (Figure 2D, 10 mM: 52.78 ± 7.24% cell loss; 15 mM: 86.47 ± 3.90% cell loss; 20 mM: 88.54 ± 1.66% cell loss; 10 mM vs 15 mM and vs 20 mM p<0.001; n=4/6/5 animals), whereas *Gad67* mRNA^+^ interneurons were equally lost in all groups (Figure 2E, 10 mM: 78.33 ± 3.77% cell loss; 15 mM: 93.35 ± 0.71% cell loss; 20 mM: 85.20 ± 4.35% cell loss; n=4/3/5 animals). Animals with a greater loss of NeuN^+^ neurons in the hilus also had a larger volume of the dispersed GCL (Figure 2H).

Next, we investigated the characteristics of epileptiform activity in chronically epileptic mice using LFP recordings at three positions in the ipsi- and contralateral hippocampus (idHC, ivHC and cdHC) (Figure 3). Interestingly, epileptic bursts occurred in a region-specific manner depending on the KA concentration: A low KA concentration (10 mM), associated with mild hippocampal sclerosis, resulted in a spatially more restricted pattern of epileptic bursts compared to high concentrations (15 and 20 mM), associated with strong hippocampal sclerosis and more widespread bursts (Figure 3A, B, I). A detailed analysis of seizure classes (according to three categories ‘high-’, ‘medium-’ and ‘low-load’ bursts) demonstrated that 10 mM KA resulted in fewer high-load events compared to 15 or 20 mM KA (Figure 3C, D, individual values in Source Data Table 1). In addition, we found that the overall mean burst ratio and epileptic spike rate were smaller for the low KA concentration with mild sclerosis (Figure 3E, burst ratios: 10 mM: 0.06 ± 0.01; 15 mM: 0.20 ± 0.03; 20 mM: 0.19 ± 0.02, 10 mM vs 15 mM and vs 20 mM p<0.01; and Figure 3F, spike rates: 10 mM: 0.32 ± 0.02; 15 mM: 0.77 ± 0.11; 20 mM: 0.71 ± 0.07 Hz, 10 mM vs 15 mM p<0.01 and vs 20 mM p<0.05; n=4/6/5 animals). Taken together, the extent of GCD and hilar cell loss was positively correlated with the spontaneous emergence of high-load events (Figure 3G, p<0.001, Pearson’s r=0.92; and Figure 3H, p<0.05, Pearson’s r=0.62; both n=15 animals).

**Figure 3.**
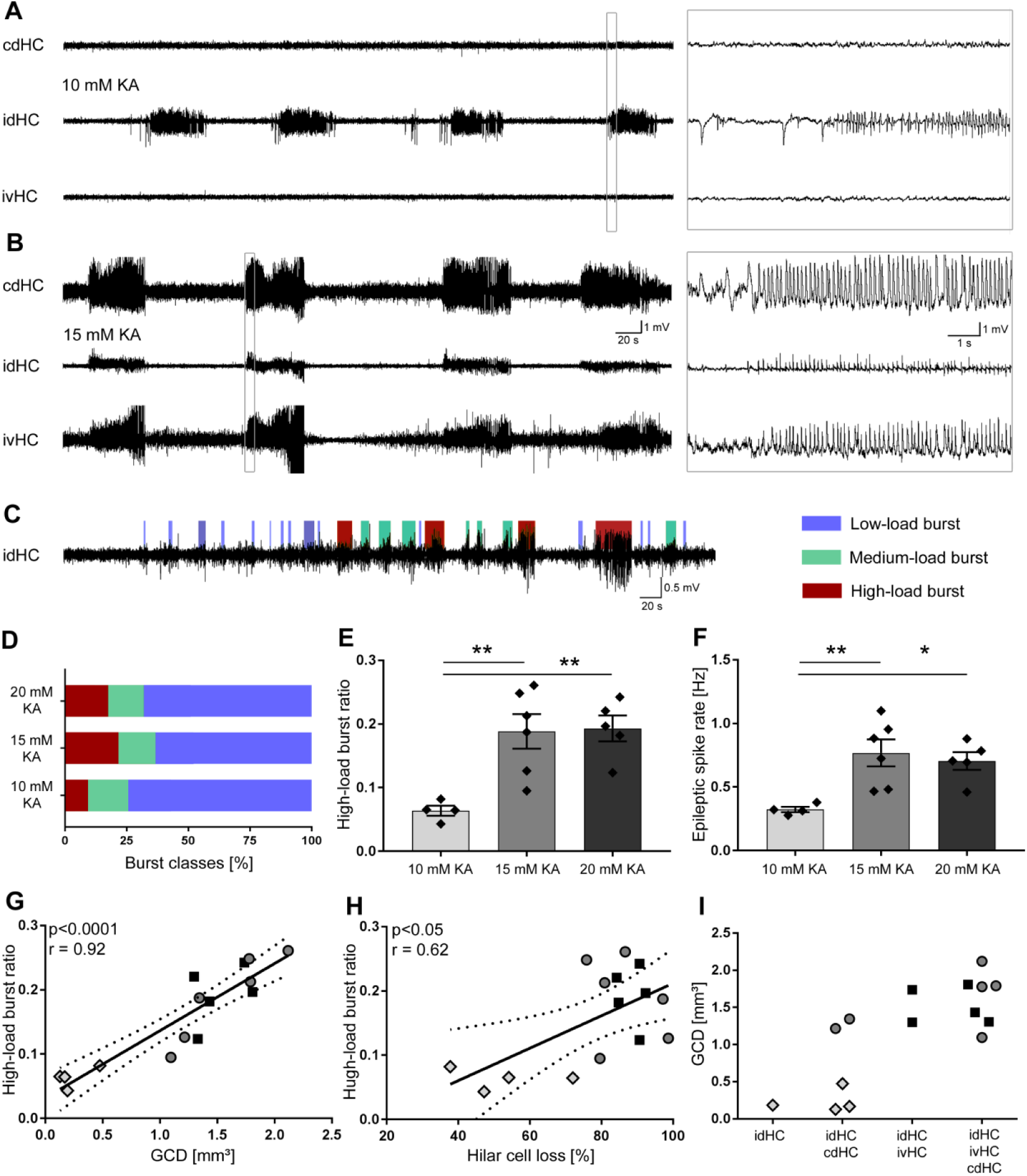
Severity and spatial occurrence of epileptiform activity elicited by different KA concentrations. **(A, B)** Representative LFP traces for low (10 mM) and high (15 mM) KA concentrations (20 mM not shown) at the three different recording sites: cdHC, idHC and ivHC. Animals with a high KA concentration show a more widespread epileptiform activity pattern. **(C – F)** Automatic quantification of epileptiform activity. High KA concentrations elicit a higher percentage of high-load burst and increased epileptic spike rate. One-way ANOVA; Tukey’s multiple comparison test; *p<0.05, **p<0.01. All values are given as mean ± SEM. **(G, H)** The burst ratio is positively correlated with GCD and hilar NeuN^+^ cell loss (**(G)**: p<0.001, two-tailed; Pearson’s r = 0.92; **(H)**: p<0.05, two-tailed; Pearson’s r = 0.62). **(I)** Pattern of epileptiform activity (visible bursts at recording sites) according to KA concentration and GCD. The epileptiform activity pattern appears spatially more restricted to the dorsal regions of the hippocampus for the low KA concentration (10 mM, shown in **A**) accompanied by a smaller extent of GCD.

Our results show that the disease severity on both, the anatomical and electrophysiological level was modulated by the strength of the excitotoxic insult, thus providing a valuable framework for the robustness of our oLFS experiments.

### oLFS of granule cells prevents spontaneous recurrent seizures

Since the sclerotic hippocampus is regarded as the epileptic focus (Esther Krook-Magnuson et al., 2015; Pallud et al., 2011), we decided to target surviving DGCs by photostimulation of afferent entorhinal fibers for seizure interference. To this end, adult mice received intrahippocampal KA and a ChR2-carrying viral construct into the medial entorhinal cortex followed by LFP recordings and oLFS in the chronic epileptic phase (Figure 4A). Prior to photostimulation, baseline activity was recorded in ‘pre’ sessions for one hour to confirm the occurrence of recurrent seizures at the idHC position (Figure 4B). We stimulated ChR2-expressing entorhinal fibers locally in the middle molecular layer of the sclerotic hippocampus (Figure 4C) with three frequencies (1, 0.5 and 0.2 Hz) in all epileptic animals that displayed different degrees of hippocampal sclerosis.

**Figure 4.**
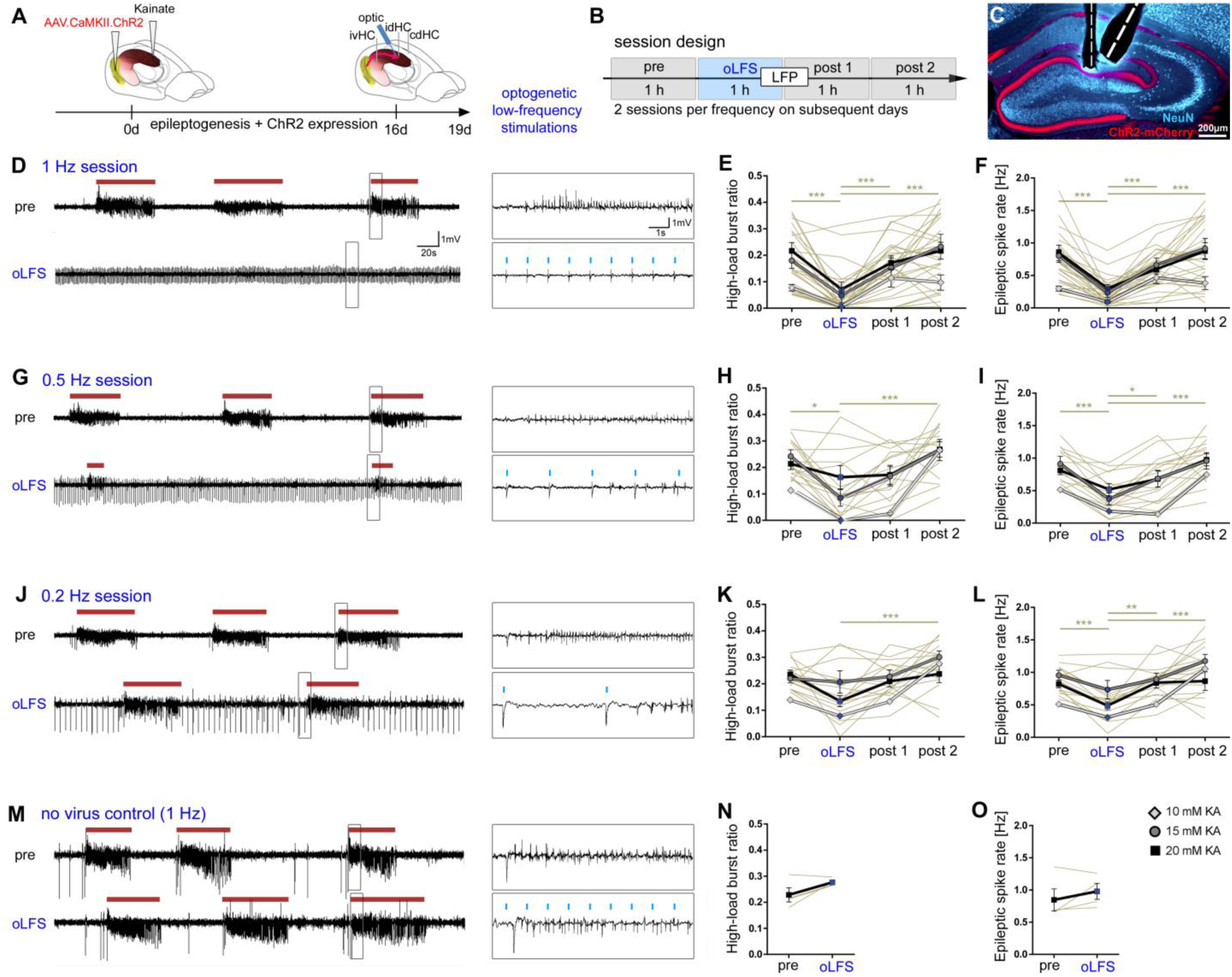
Optogenetic low-frequency stimulation of entorhinal afferents interferes with ictogenesis. **(A – C)** Experimental design. We targeted ChR2 expression (**C**, red) to excitatory neurons in the medial entorhinal cortex using viral vectors and **(B, C)** locally stimulated entorhinal afferents in the sclerotic idHC for one hour per day, twice at each frequency applying only one frequency per session (1, 0.5 or 0.2 Hz). **(D, G, J, M)** Representative LFP traces (15 mM KA, idHC electrode) for the ‘pre’ and ‘oLFS’ sub-sessions (1, 0.5, 0.2 Hz and no virus control, 1 Hz) and are shown. **(D)** Optogenetic stimulation with pulsed light at 1 Hz effectively decreases spontaneous epileptiform activity (marked with a red bar). **(E, F)** Automatic quantification of epileptiform activity shows that oLFS reduces the burst ratio as well as the epileptic spike rate in all animals independently of the KA concentration (10 mM: light grey; 15 mM: dark grey; 20 mM: black) followed by a return to pre-stimulation levels within two hours (‘post 1’ and ‘post 2’. **(G, J)** oLFS with 0.5 Hz or 0.2 Hz has a weaker antiepileptic effect as quantified with the automatic detection algorithm **(H, I and K, L).** Single sessions (beige) were used to calculate the RM one-way ANOVA; Tukey’s multiple comparison test (all KA concentrations merged); *p<0.05, **p<0.01 and ***p<0.001. **(M – O)** 1 Hz stimulation doesn’t have an effect in no-virus controls (20 mM KA). All values are given as mean ± SEM.

One hour of optogenetic stimulation with pulsed light at 1 Hz effectively suppressed the epileptiform activity (burst ratio) and reduced the epileptic spike rate in almost all animals independent of the KA concentration. When oLFS was stopped, epileptic activity returned to pre-stimulation levels within two hours (Figure 4D – F, individual values in Source Data Table 2). Photostimulation at both 0.5 and 0.2 Hz had also suppressive effects on epileptiform activity, but less pronounced than 1 Hz (Figure 4G – L, values in Source Data Table 2). oLFS (1 Hz) did not influence epileptiform activity in no-virus control mice (Figure 4M – O, individual values in Source Data Table 2).

To clarify whether the anti-epileptic effect was locally restricted to the stimulation site (idHC), we analyzed LFPs at the other two electrode positions (ivHC and cdHC). Interestingly, epileptiform activity was also suppressed at these sites (data not shown) indicating that oLFS in the sclerotic focus was highly effective in the whole hippocampus.

In parallel to LFP recordings and optogenetic stimulation, we assessed if the animals’ motor behavior in the open field was influenced by oLFS. We observed that mice showed frequent grooming and exploration during photostimulation. Video tracking revealed that all mice, independent of the KA concentration, exhibited normal running behavior that declined gradually during the recording time of four hours (Supplementary Figure 2A – D, individual values in Source Data Table 3). Analysis of all ‘pre’ and ‘oLFS’ sessions showed a stable pattern over the six days of stimulation (Supplementary Figure 2E, F), indicating that hippocampal oLFS did not impair open-field behavior of chronically epileptic mice.

Taken together, oLFS of entorhinal afferents at 1 Hz was most effective compared to lower frequencies in preventing the emergence of spontaneous recurrent seizures during stimulation independently of the disease severity.

### Analysis of evoked responses to oLFS

Since oLFS was not equally effective in suppressing epileptiform activity in all sessions (see Figure 4), we analyzed the strength of the evoked responses to photostimulation by determining the median AUC in each session for all frequencies and electrode positions (see Methods).

Linear regression analysis revealed a positive relationship between the median evoked response and the stimulation efficacy (quotient of the burst ratio of ‘oLFS’ and ‘pre’ sub-sessions) especially for 1 and 0.5 Hz (Supplementary Figure 3A – C) indicating that a strong cellular response is necessary for successful seizure suppression. This observation was most pronounced in the sclerotic focus where we applied photostimulation. When we compared the efficacy of stimulation across frequencies, we considered only sessions with a cellular response within the range of the standard deviation of the mean. Thus, oLFS at 1 Hz resulted in a remarkable reduction (90%) of the burst ratio, whereas 0.5 Hz and 0.2 Hz were less efficient (80% and 40% reduction), but nevertheless showed anti-epileptic effects (Supplementary Figure 3D, 1 Hz: 92.17%; 0.5 Hz: 78.93%; 0.2 Hz: 39.42%, 95% CI [80.49, 97.86], [44.32, 100.00], and [18.11, 50.42], respectively, 0.2 Hz vs 1 Hz p<0.001 and vs 0.5 Hz p<0.05; n=29/18/17 sessions; 15/12/12 animals).

Next, we tested whether the neuronal response to oLFS was confined to the stimulated area. To this end, we analyzed the spatial and temporal occurrence of evoked responses at all three recording sites. In all animals, light-pulses delivered to the idHC did not only trigger local but also delayed responses in both hippocampi (Supplementary Figure 4). Population spikes occurred first at the stimulation site in the sclerotic region followed by the ipsilateral ventral and the contralateral dorsal position. These latencies remained stable over the stimulation period of one hour (Supplementary Figure 4D, idHC to ivHC: 3.96 ± 0.30 ms; idHC to cdHC: 8.99 ± 0.59 ms, n=13/8 sessions), suggesting that photostimulation of entorhinal fibers may lead to action potential generation in a subset of DGCs and subsequent propagation within the hippocampal network.

To infer if anti-ictogenic effects by oLFS are due to activation of DGCs or are specific for the stimulation of entorhinal afferents, we performed direct photostimulation of ChR2-expressing DGCs (Figure 5A, B). Similar to stimulation of entorhinal afferents, 1 Hz oLFS of DGCs achieved substantial seizure control (reduction of burst ratio and epileptic spike rate) with no apparent rebound effect after the oLFS offset (Figure 5C – E, individual values in Source Data Table 4, n=3 animals). However, in contrast to perforant path oLFS, the reappearance of high-load bursts was evident within minutes after direct photostimulation of DGCs (Figure 5F, 1 Hz perforant path oLFS: 35.20 ± 7.14 min; 1 Hz DGC oLFS: 7.93 ± 2.84 min, p<0.001; n=18/7 sessions; 11/3 animals).

**Figure 5.**
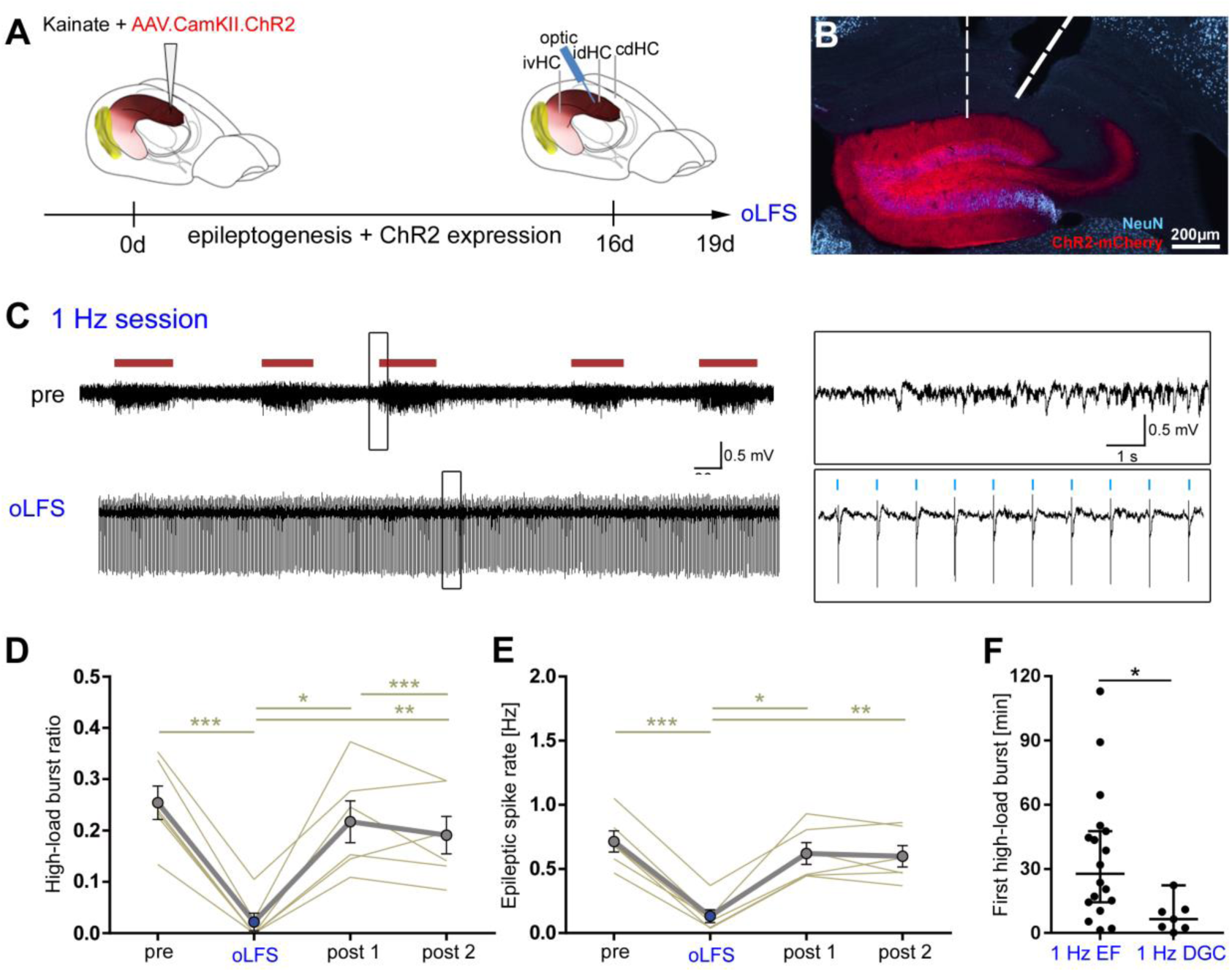
Direct oLFS of DGCs results in suppression of epileptiform activity with a shorter seizure-free period after stimulation. **(A, B)** Experimental design. To infer if anti-ictogenic stimulation effects were due to activation of DGCs or specific to stimulation of entorhinal afferents, we performed direct photostimulation of ChR2-expressing DGCs in idHC (**B**, red). Animals were injected with KA (15 mM) and virus in the dentate gyrus. **(C)** An exemplary LFP trace from the idHC site. **(D, E)** 1 Hz stimulation of DGCs significantly reduces the high-load burst ratio and epileptic spike rate. Epileptiform activity reoccurs within the first hour after stimulation (‘post 1’) and remains in the second hour (‘post 2’). RM one-way ANOVA; Tukey’s multiple comparison test; *p<0.05, **p<0.01 and ***p<0.001. Values are given as mean ± SEM. **(F)** Compared to entorhinal fiber (EF) stimulation, seizure-activity returned directly after stimulation cessation. Unpaired t-test; *p<0.05.

In conclusion, both, direct oLFS of DGCs and indirect oLFS via entorhinal afferents had seizure suppressive effects, with the latter showing a longer-lasting anti-epileptic effect of around 30 min.

### oLFS interferes with seizure generalization

Since spontaneous recurrent seizures are mainly subclinical in the intrahippocampal KA mouse model and seizure generalization is rare, we aimed at inducing generalized seizures by photostimulation (Janz et al., 2018; Osawa, Iwasaki, Hosaka, Matsuzaka, & Tomita, 2013). Initial experiments with 10 Hz for 10 s showed that during the first few seconds of stimulation, only evoked potentials followed each light pulse. High-amplitude epileptic spikes emerged in addition to the evoked potentials during further stimulation and gradually became rhythmic and dominant, progressing into a fully-blown behavioral seizure (Supplementary Figure 5A). On the electrophysiological level, these seizures displayed electrographic features highly similar to those of spontaneous generalized seizures (Supplementary Figure 5B) and were also accompanied by the same stereotypic myoclonic movements (e.g., rearing, falling and convulsion).

First, we determined the minimum stimulus duration sufficient to reliably trigger a generalized seizure but avoiding that the pro-convulsive 10 Hz stimulus masks the anti-ictogenic effect of oLFS (Figure 6A). Interestingly, in mice with lower KA concentrations generalized seizures were induced much faster, suggesting a higher susceptibility for seizure generalization (Figure 6B, 10 mM: 5.75 ± 0.63 s; 15 mM: 7.83 ± 0.87 s; 20 mM: 13.67 ± 1.86 s, 10 and 15 mM vs 20 mM p<0.01; n=3/6/4 animals). With the progression of seizure activity, mice exhibited behavioral symptoms equivalent to RS stages 1 to 5 independent of the stimulation duration (Figure 6C, n=3/6/4 animals).

**Figure 6.**
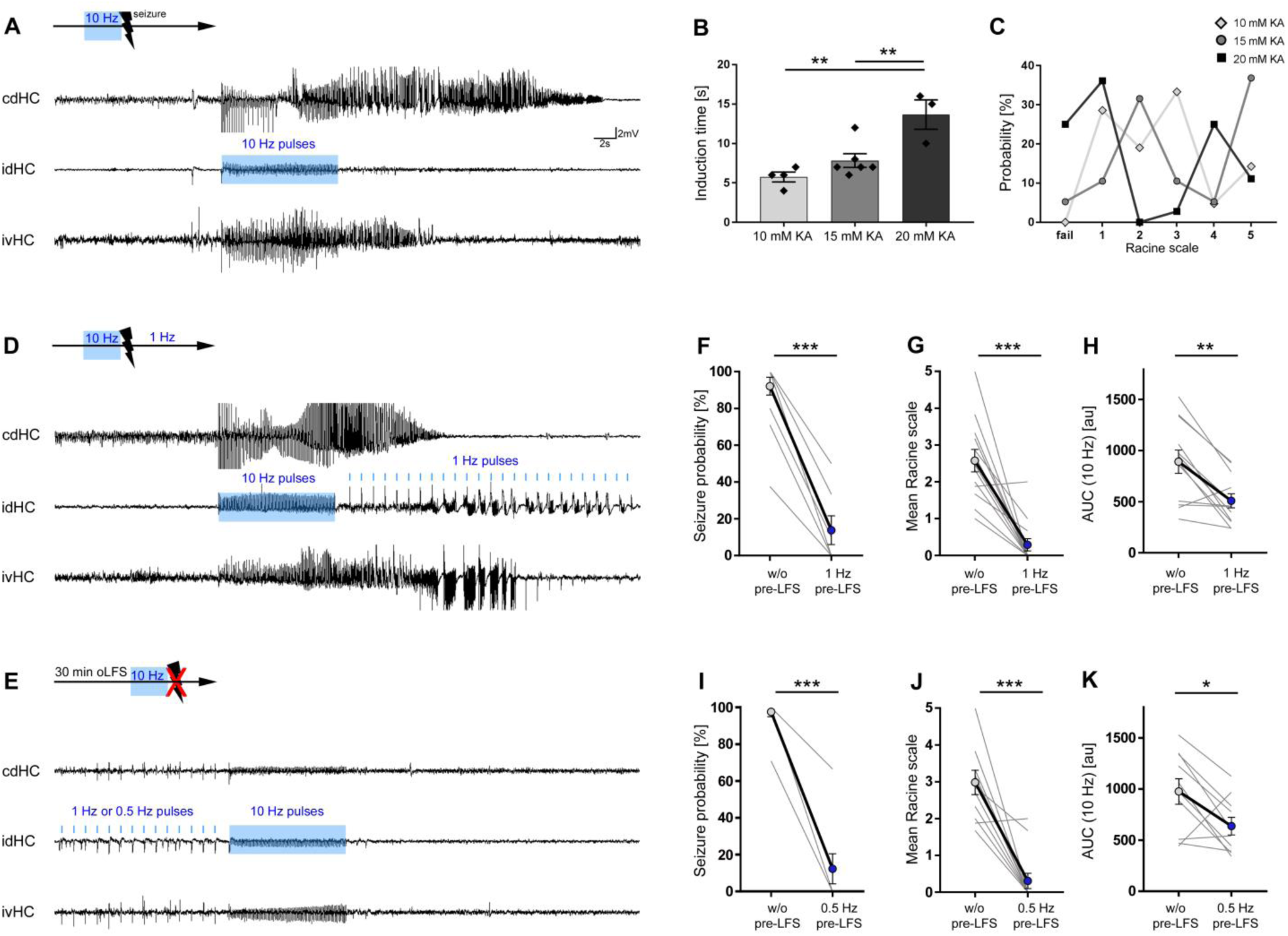
Preceding oLFS prevents seizure generalization. **(A, D, E)** Representative LFP traces at the three recording sites (cdHC, idHC and ivHC). Schematic of the respective stimulation procedure is shown above each cdHC trace. **(A, B)** Local 10 Hz photostimulation of entorhinal afferents reliably induces generalized seizures in all KA groups. The induction time for generalized seizures depends on the KA concentration. One-way ANOVA; Tukey’s multiple comparison test; **p<0.01. **(C)** Mice exhibit behavioral symptoms equivalent to RS stage 1 to 5, independently of the KA concentration. **(D)** Seizure generalization cannot be prevented by oLFS (1 Hz) applied directly after seizure induction. **(E)** oLFS for 30 min prior to the pro-convulsive stimulus effectively prevents seizure generalization. **(F, I)** Both, 1 and 0.5 Hz oLFS significantly decrease the seizure probability in all animals. Wilcoxon rank test, matched-pairs; ***p<0.001 (n=14 animals). **(G, J)** Trials in which seizure generalization has not been prevented completely, the ensuing seizure is associated with a milder behavioral phenotype (RS stage). Paired t-test; ***p<0.001. **(H**, **K)** Cellular response to 10 Hz stimulation quantified as mean AUC. The response is reduced after 1 or 0.5 Hz oLFS stimulation in sessions in which seizures have been successfully suppressed. Paired t-test; **p<0.01 and *p<0.05 (n=13 and 10 animals), respectively. All values are given as mean ± SEM.

Next, we probed the competence of oLFS to interfere with generalized seizures. When we started 1 Hz oLFS directly after the pro-convulsive 10 Hz stimulus, we could not interrupt ongoing seizures (Figure 6D). In contrast, pre-conditioning with oLFS for 30 minutes applied prior to the pro-convulsive stimulus at either 1 or 0.5 Hz very effectively lowered the probability for seizure generalization (Figure 6E, F, I, without (w/o) pre-oLFS: 96.79 ± 2.20%; with 1 Hz pre-oLFS: 15.60 ± 7.75%, p<0.01; n=7; w/o pre-oLFS: 97.73 ± 2.27%; with 0.5 Hz pre-oLFS: 12.12 ± 8.13%, p<0.05; n=14/11 animals). Those trials in which seizure generalization was not prevented completely, the ensuing seizure was associated with a milder behavioral phenotype (Figure 6G, w/o oLFS: RS 2.58 ± 0.31; with 1 Hz oLFS: RS 0.29 ± 0.17; Figure 6J, w/o oLFS: RS 2.88 ± 0.32; with 0.5 Hz oLFS: RS 0.37 ± 0.25, p<0.001; n=13/10 animals).

In conclusion, oLFS of entorhinal afferents not only prevents subclinical, spontaneous recurrent seizures but also interferes with the generation of evoked generalized seizures.

### Effects of oLFS on the cellular response

In order to understand the underlying mechanisms of the anti-ictogenic effects of oLFS, we investigated the evoked responses in DGCs by quantifying the AUC of each individual evoked response over the one hour stimulation period. For this analysis, we chose only sessions, which were within the 95% CI regarding the respective stimulation efficacy, as reported above (Supplementary Figure 3D).

Both, indirect, via entorhinal afferents, or direct oLFS of DGCs, evoked stable responses with respect to temporal occurrence and waveforms (Figure 7). Responses evoked by direct photostimulation of DGCs decreased slightly over time (Figure 7A1-3, B), whereas stimulation of entorhinal afferents caused a more pronounced and rapid (within 10 min) decrease of the cellular response (Figure 7C1-3, D) suggesting that a synaptic mechanism contributes to the longer-lasting anti-ictogenic effect of entorhinal afferent-mediated oLFS. Lower frequencies (0.5 and 0.2 Hz) altered the cellular response much less (Figure 7E – H). Similarly, pre-conditioning with 1 or 0.5 Hz reduced the evoked response (AUC) of the pro-convulsive 10 Hz pulse-train by about 40% (Figure 6H, w/o oLFS: 890.9 ± 113.4; with 1 Hz oLFS: 508.6 ± 68.57, p<0.01; Figure 6K, w/o oLFS: 977.6 ± 125.4; with 0.5 Hz oLFS: 638.8 ± 86.05, p<0.05, n=12/10 animals).

**Figure 7.**
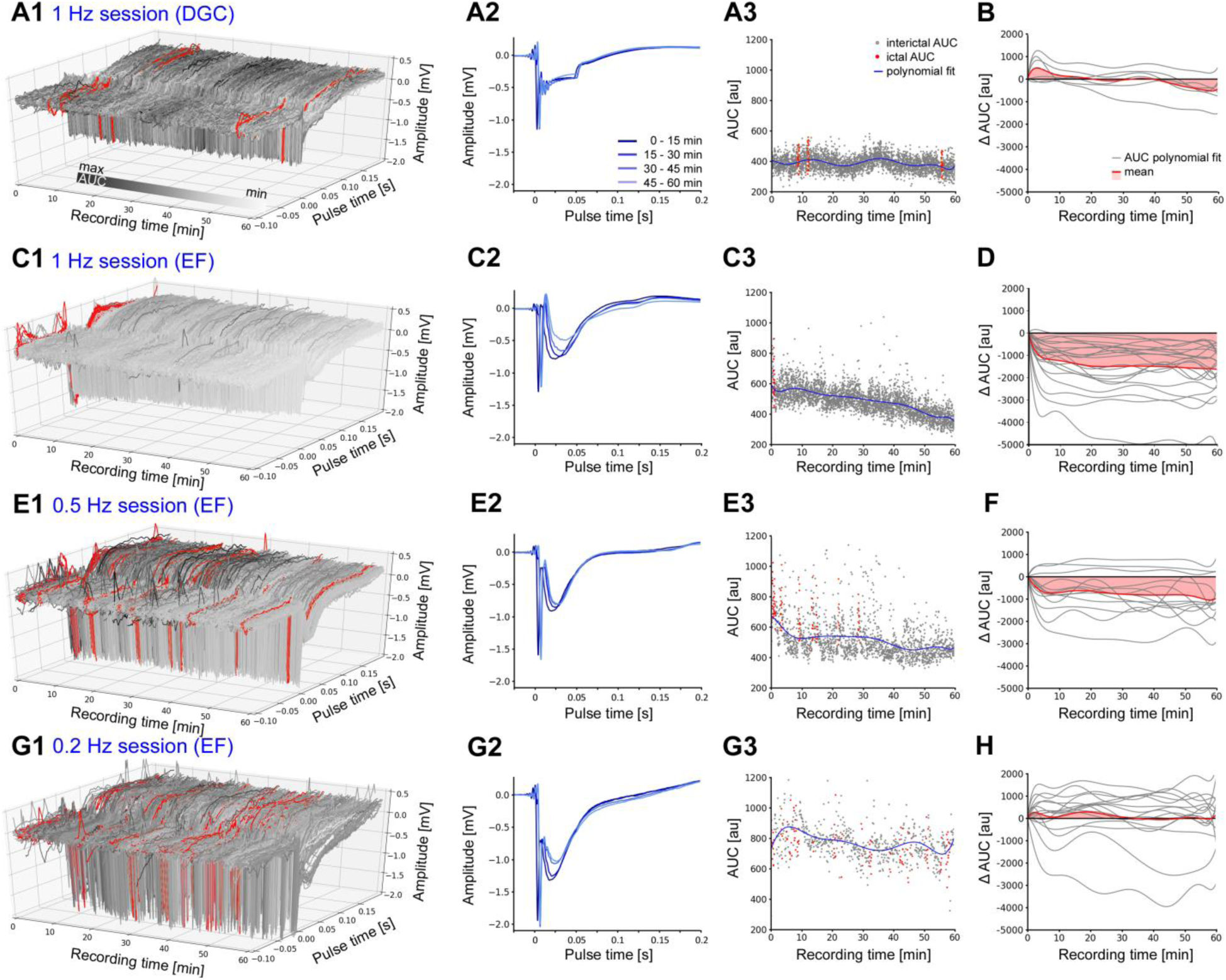
Evoked cellular responses decrease over time during continuous oLFS. (**A1)** Representative examples of evoked responses in DGCs following direct or **(C1, E1, G1)** indirect photostimulation via entorhinal fiber (EF) stimulation at different frequencies. **(A2, C2, E2, G2)** Mean evoked response for 15 min time windows are shown. **(A3, C3, E3, G3)** For each evoked response, magnitudes are measured using the AUC during a peri-stimulus time interval [-0.1, +0.2 s] upon each light pulse. Responses during ictal periods (high-load events) are marked in red. A polynomial fit of single interictal AUC values is shown (blue line). **(A1)** Direct photostimulation of DGCs (1 Hz) evokes constant responses typical for ChR2 activation. **(A2)** Evoked cellular responses and **(A3)** single AUC values are stable over time. **(B, D, F, H)** Normalized polynomial fits (delta AUC) for all stimulation sessions (grey) and mean changes (red). **(B)** Evoked responses stay stable over one hour direct photostimulation. **(C1-3)** Entorhinal fiber stimulation (1 Hz) causes a decline of the cellular response over time. **(D)** Mean evoked responses (AUCs) decrease strongly within the first 10 min of photostimulation. **(E – H)** Entorhinal fiber stimulation at 0.5 and 0.2 Hz has a weaker effect on the change of the evoked response magnitude.

To elaborate whether oLFS decreases the excitability of DGCs, we studied their intrinsic properties and the synaptic strength of entorhinal inputs in acute slices from chronically epileptic mice. Whole-cell recordings were obtained from DGCs located in the outer region of the dispersed GCL. Cells were filled with biocytin during recordings for subsequent morphological identification (Figure 8A, n=11 animals). Photostimulation of afferent entorhinal fibers (473 nm; 50 ms pulse duration) induced robust depolarization (5.1 ± 1 mV, n=14 cells) and was occasionally sufficient to induce action potentials (−70 mV holding potential, n=4 cells). During oLFS (10 min, 1 Hz) evoked synaptic responses were strongly depressed (Figure 8B, reduction of EPSP amplitude to 28.5 ± 9.9% of the original response, n=14 cells), suggesting synaptic fatigue. To test this possibility, we evaluated the effect of oLFS on discharge probability upon electrical stimulation of entorhinal fibers. To maintain intracellular conditions in DGCs, we used cell-attached recordings. Action potentials were reliably evoked in DGCs by five electrical stimulation pulses at 50 Hz. Photostimulation for 10 min, however, clearly reduced discharge probability (Figure 8C, discharge probability at 100 V stimulation, control: 35.4 ± 6.0% vs. after oLFS: 21.2 ± 5.2%, p<0.001, n=19 cells). In contrast, intrinsic properties of DGCs were not altered by the applied stimulation protocol (control: resting membrane potential (V_m_) = −73.4 ± 1.1 mV; C_m_ = 53.5 ± 3.7 pF; R_m_ = 298.7 ± 27.4 MΩ; Rheobase = 150.8 ± 14.1 pA; n=27 cells; after oLFS: V_m_ = −74.8 ± 1.3 mV; C_m_ = 56.4 ± 7.5 pF; R_m_ = 313.9 ± 36.8 MΩ; Rheobase = 153.3 ± 26.4 pA; n=14 cells) suggesting that the reduced excitability of DGCs during oLFS may be explained by a reduced glutamate release from entorhinal projections.

**Figure 8.**
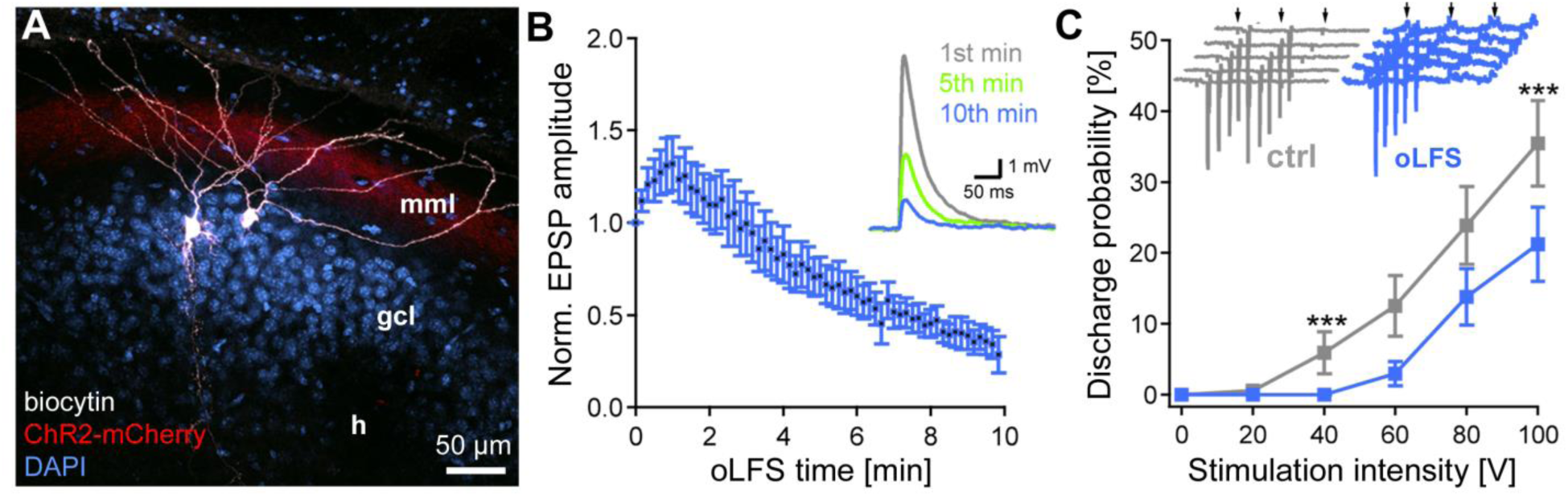
Decrease of single-cell EPSPs and discharge probability after 10 min oLFS. **(A)** Representative confocal projection of a dentate gyrus slice from a KA-(15 mM) and AAV-treated mouse 28 days after SE. Two biocytin-filled DGCs (white) have been recorded in this section. ChR2-expressing entorhinal fibers are visible in the middle molecular layer (red, mml). Cell bodies are stained with DAPI (blue). h, hilus; scale bar 50 µm. **(B)** Blue light pulses induce robust EPSPs which decline strongly during a 10 min stimulation protocol (50 ms pulses at 1 Hz). **(C)** Extracellular electrical stimulation (5 pulses, 50 Hz, arrows) of entorhinal fibers induces action potentials measured by loose-patch recordings of DGCs (inset, gray traces). Photostimulation at 1 Hz over 10 min (inset, blue traces) significantly reduces the discharge probability of DGCs. ANOVA on ranks with Dunn’s post-hoc correction; ***p<0.001. Values are given as mean ± SEM.

## Discussion

In the present study, we applied oLFS of entorhinal afferents to the epileptic hippocampus in experimental MTLE to interfere with ictogenesis *in vivo*. In particular, 1 Hz stimulation demonstrated remarkable anti-ictogenic effects which were independent of the individual degree of hippocampal sclerosis and seizure severity. Reaching out towards an understanding of the underlying mechanism, our findings suggest that oLFS in the epileptic focus lowers seizure susceptibility most likely by synaptic fatigue rather than by decreasing the intrinsic excitability of DGCs.

To date, electrical high-frequency stimulation (HFS, 130-200 Hz and 1-5 V) of the hippocampus has been used empirically in clinical settings as an approach to control intractable mesiotemporal seizures. It is widely assumed that electrical pulses applied at high frequencies reduce the seizure threshold by disrupting network synchronization, while low frequencies are believed to elicit the opposite effect. So far, only a few studies applied LFS in human MTLE (Lim et al., 2016), whereas HFS has been performed extensively with variable success (Fisher & Velasco, 2014; Li & Cook, 2018): Half of all patients from clinical studies experienced a 48–95% seizure reduction; however, in randomized controlled trials the overall efficacy reached only 26–40% following HFS (Li & Cook, 2018). This variability can be partially explained by sub-optimal electrode positioning, different follow-up durations and preselection criteria. Importantly, the extent of hippocampal sclerosis appeared to be a critical parameter for stimulation efficacy (Boëx et al., 2011; Lim et al., 2016; Velasco et al., 2007). This is in line with the hypothesis that neuronal loss and/or altered electrical resistance in sclerotic neural tissue hinder HFS, since stimulation can only be successful when targeting a sufficiently preserved network. Therefore, patients with hippocampal sclerosis may require specific stimulation parameters to achieve seizure control. In fact, a small cohort study (Lim et al., 2016) showed that LFS at 5 Hz was effective in two patients with hippocampal sclerosis pointing to the use of LFS in a clinical setting. However, to systematically assess anti-ictogenic effects of LFS in relation to disease parameters, studies in translational animal models are crucial.

Accordingly, we used the intrahippocampal KA mouse model of MTLE that recapitulates the major pathological hallmarks of the human disease, comprising unilateral hippocampal sclerosis and the emergence of spontaneous recurrent seizures (Bouilleret et al., 1999; Janz et al., 2017b). An important aspect of our study was to create an animal model that reflects the inter-individual variability of neurodegeneration and seizure frequency as observed in human MTLE (Blümcke et al., 1999; Thom, 2014). In fact, we found experimental conditions that resulted in varying degrees of histopathological changes and of epileptiform activity, depending on the KA concentration. In particular, cell death of hilar neurons and the extent of GCD increased with the KA dose, whereas DGCs survived. This is in line with previous reports from both, human patients and mouse epilepsy models, showing that DGCs are highly resistant to overexcitation and are the surviving neuronal sup-type under sclerotic conditions (Thom, 2014; Young et al., 2009). Therefore, we chose DGCs as targets and photostimulated entorhinal fibers which synapse onto DGCs dendrites, thus stimulating DGCs indirectly. In fact, oLFS of entorhinal afferents at 1 Hz effectively abolished spontaneous recurrent seizures in animals with both mild and severe hippocampal sclerosis. Also lower stimulation frequencies (0.5 and 0.2 Hz) showed reasonable anti-epileptic effects. In addition, this protocol appeared to generate action potentials in DGCs, since evoked populations spikes were not only evident at the site of light delivery, but also with a polysynaptic delay in the other ipsilateral and contralateral recording sites, showing that the neuronal network necessary for propagation was maintained. Accordingly, local oLFS effectively suppressed seizure activity in both hippocampi.

Although the mechanism underlying the anti-epileptic effect of oLFS is not clear, several lines of evidence suggest that LFS, either electrically or optogenetically, reduces the excitability of the hippocampus and associated networks (i.e., the entorhinal cortex). *In vitro* data indicate that electrical stimulation of the subiculum or entorhinal cortex at 1 Hz suppresses ictal activity induced by 4-AP (Shiri et al., 2017). Similarly, 1 Hz photostimulation of CaMKIIa-positive principal cells in the entorhinal cortex reduces the frequency and duration of ictal discharges (Shiri et al., 2017). Also, 1 Hz electrical or optogenetic stimulation of the entorhinal cortex *in vivo* successfully reduces the severity of seizure-like after-discharges upon hippocampal kindling in mice (Xu et al., 2016). Similar results were obtained by direct electrical LFS in the ventral hippocampal fissure in both, a genetic and a kindling epilepsy model (Kile, Tian, & Durand, 2010; Rashid, Pho, Czigler, Werz, & Durand, 2012). Although these studies provide valuable insight into the seizure-suppressing effects of LFS, it was unclear whether LFS is capable of preventing ictogenesis in chronic MTLE with hippocampal sclerosis. In this context, our study reveals an unprecedented efficacy of 1 Hz stimulation directly in the highly-sclerotic hippocampus to interfere with the generation of both, spontaneous subclinical recurrent seizures and evoked generalized seizures. In contrast to Xu et al. (2016) (Xu et al., 2016), our results suggest that the anti-ictogenic effect does not depend on the local recruitment of GABAergic interneurons since this cell population is extensively diminished in our model (Marx, Haas, & Häussler, 2013). This is an important observation because the loss of GABAergic interneurons is also a common finding in human MTLE with hippocampal sclerosis (Young et al., 2009). Therefore, targeting GABAergic interneurons is not feasible in these cases. The majority of current optogenetic approaches for seizure control in experimental epilepsy rely on decreasing network activity, either by direct inhibition of DGCs or by activating GABAergic interneurons (Kokaia et al., 2013; Esther Krook-Magnuson et al., 2013; Ladas et al., 2015; Ledri et al., 2014), whereas our approach aims at low-frequency activation of ‘epileptic’ DGCs by local stimulation of entorhinal afferents. Considering the clinical limitations of optogenetics, our approach could nevertheless be translated into the clinics, as from an anatomical perspective, selective stimulation of entorhinal afferents in humans can be achieved by electrode placement in the clearly-defined angular bundle (Zeineh et al., 2017). Therefore, indirect stimulation via entorhinal afferents represents a highly promising strategy that may overcome limited stimulation efficacy associated with hippocampal sclerosis.

Mechanistically, it is rather astonishing that synchronization of the hippocampal network by oLFS via entorhinal afferents interferes with ictogenesis. A hypothesis put forward by Avoli et al. (2013) (Avoli, de Curtis, & Köhling, 2013) implies that more frequent activation (i.e. at 0.5 Hz vs. 0.1 Hz) of the network results in less accumulation of extracellular potassium compared to large potassium efflux associated with GABA_A_ receptor-mediated interictal discharges that could trigger a seizure. Clamping GABA-mediated potentials with 1 Hz LFS (directly via activation of GABAergic interneurons or indirectly via feedback inhibition upon principal cell activity) could therefore restrain pro-ictogenic discharges (Shiri et al., 2017). Our results show that this intriguing mechanism cannot fully account for the anti-ictogenic effect of oLFS, since in whole-cell recordings, we observed a decrease in the amplitude of evoked responses in the presence of GABA_A_ and GABA_B_ receptor blockers. Another explanation, supported by our data, relies on synaptic depression (both, synaptic fatigue and long-term depression (LTD)) that is induced at the entorhinal–DGC synapse with stimulation frequencies equal or below 1 Hz (Abrahamsson, Gustafsson, & Hanse, 2005; Gonzalez, Morales, Villarreal, & Derrick, 2014). Accordingly, we show that the amplitude of evoked responses is decreased over time on both, the network and the single-cell level, which could explain the delayed onset of spontaneous seizures following oLFS. Although not tested in the present study, we propose that our oLFS paradigm leads to hetero-rather than homosynaptic depression, since it is unlikely that downscaling of entorhinal inputs only is sufficient to suppress seizure activity given that performant path transection does not alleviate seizures (Meyer, Kienzler-Norwood, Bauer, Rosenow, & Norwood, 2016). Yet, synaptic depression may only account for the prolonged anti-ictogenic effects observed upon oLFS of entorhinal afferents, in which the gradual decrease of evoked potentials was much more prominent compared to direct oLFS of DGCs, although, both protocols showed remarkable performance in acute seizure suppression. Therefore, with respect to acute anti-ictogenic effects, we suggest that oLFS drives the hippocampal network into a “stable state”, reducing the probability of seizures while LFS is ongoing. This notion is supported by Chang et al., (2018) (Chang et al., 2018) who showed that evoked interictal-like discharges elicit anti-ictogenic effects that depend on both, the network state and stimulus properties, including amplitude and frequency. This study investigated the influence of interictal discharges on ictogenesis *in vitro* in local CA3 and CA1 circuits, two subnetworks that are typically lost in MTLE with hippocampal sclerosis. Our results strongly suggest that the authors’ observations also apply to the entorhinal-dentate network *in vivo*. In line with this interpretation, oLFS of entorhinal afferents may introduce glutamatergic synaptic perturbation, which is followed by transient suppression of neuronal activity (M. De Curtis, Librizzi, & Biella, 2001; Marco De Curtis & Avanzini, 2001; Muldoon et al., 2015) and interference with the slow process of critical slowing before seizure onset (Chang et al., 2018).

In conclusion, our study has identified 1 Hz oLFS of entorhinal afferents as a highly efficient approach to interfere with ictogenesis in MTLE with hippocampal sclerosis. We show that the effect is largely driven by repetitive activation of DGCs residing in the seizure focus, and we have shed light on the associated cellular mechanisms. Considering the potential for clinical translation, our findings may pave the way for effective seizure control in one of the most common forms of drug-resistant epilepsy.

## Materials and Methods

### Animals

Experiments were conducted with adult (8–12 weeks) transgenic male mice (C57BL/6-Tg(Thy1-eGFP)-M-Line) (Feng et al., 2000). Each animal represents an individual experiment, performed once. A total of 53 mice were used for this study. Mice were kept in a 12 h light/dark cycle at room temperature (RT) with food and water *ad libitum*. All animal procedures were carried out in accordance with the guidelines of the European Community’s Council Directive of 22 September 2010 (2010/63/EU) and were approved by the regional council (Regierungspräsidium Freiburg).

### KA and virus injections

Mice were injected with KA into the right dorsal hippocampus (n=33 for in vivo oLFS experiments and n=20 for acute slice electrophysiology), as described previously (Heinrich et al., 2006; Häussler et al., 2012; Janz et al., 2017a). Accordingly, mice were deeply anesthetized (ketamine hydrochloride 100 mg/kg, xylazine 5 mg/kg, atropine 0.1 mg/kg body weight, i.p.) followed by stereotaxic injection of 50 nL of either 10, 15 or 20 mM KA solution (Tocris, Bristol, UK) in 0.9% sterile saline. Mice were randomly assigned to the respective kainate concentration group. Stereotaxic coordinates relative to bregma were: anterioposterior (AP) = −2.0 mm, mediolateral (ML) = −1.5 mm, and relative to the cortical surface: dorsoventral (DV) = −1.5 mm. Following KA injection the occurrence of a behavioral SE was verified, characterized by mild convulsion, chewing, immobility or rotations, as described before (Riban et al., 2002; Tulke, Haas, & Häussler, 2019). Mice which did not experience a SE (n=4) or did not survive KA treatment (n=6) were excluded from further experiments.

For optogenetic stimulation of entorhinal fibers, KA-treated animals were stereotaxically injected with a recombinant adeno-associated virus (AAV, 0.45 µl; n=22 for in vivo oLFS experiments and n=17 for acute slice electrophysiology), carrying the genomic sequences for channelrhodopsin 2 (ChR2) fused to mCherry under the control of the Ca^2+^/calmodulin-dependent kinase II alpha (CaMKIIa) promoter (AAV1.CaMKIIa.hChR2(H134R)-mCherry.WPRE.hGH; Penn Vector Core, Pennsylvania, USA) into the medial entorhinal cortex in the same surgery (Janz et al., 2018). Stereotaxic coordinates relative to bregma: AP = −5.0 mm, ML = −2.9 mm, and relative to the cortical surface: DV = −1.8 mm. KA-injected mice without virus injection were used as controls (no-virus controls, n=3). In a subset of animals, the viral vector (0.35 µl) was injected at the same location as KA to enable direct DGC stimulation (n=6).

### Implantations

Teflon-coated platinum-iridium wires (125 µm diameter; World Precision Instruments, Sarasota, Florida, USA) were implanted at three positions into the hippocampal formation in KA/virus-injected mice at 16 to 19 days after SE as described previously (Janz et al., 2018): ipsilateral dorsal (idHC), ipsilateral ventral (ivHC) and contralateral dorsal (cdHC). All animals were additionally implanted with an optic fiber (ferrule 1.25 mm, cannula 200 µm diameters; Prizmatix Ltd., Givat-Shmuel, Israel) at the same position as the idHC electrode, but at a 30° angle. Stereotaxic coordinates were chosen relative to bregma (AP, ML) or to the cortical surface (DV): cdHC: AP = −2.0 mm, ML = +1.4 mm, DV = −1.6 mm; idHC: AP = −2.0 mm, ML = −1.4 mm (−2.4 mm for the optic fiber), DV = −1.6 mm; and ivHC: AP = −3.4 mm, ML = −2.8 mm, DV = −2.1 mm. The correct positions of electrodes and optic fibers were confirmed by histology (Supplementary Figure 1). Two stainless steel screws (DIN 84; Schrauben-Jäger, Landsberg, GER) were implanted above the frontal cortex to provide a reference and ground, respectively. Electrodes and screws were soldered to a micro-connector (BLR1-type). The implant was fixed with dental cement (Paladur).

### In vivo oLFS experiments

After recovery from implantations, freely-behaving mice were first recorded on two successive days (three hours each) to determine reference LFPs (Figure 1). Each mouse represents the biological and the number of recordings per mouse the technical replicate. To this end, mice were connected to a miniature preamplifier (MPA8i, Smart Ephys/ Multi Channel Systems, Reutlingen, GER). Signals were amplified 1000-fold, bandpass-filtered from 1 Hz to 5 kHz and digitized with a sampling rate of 10 kHz (Power1401 analog-to-digital converter, Spike2 software, Cambridge Electronic Design, Cambridge, UK). On days 19 to 25 post-injection, photostimulation with pulsed blue light (460 nm; 50 ms pulse duration; 150 mW/mm² at the fiber tip; blue LED, Prizmatix Ltd.) was applied at low frequencies (1, 0.5 or 0.2 Hz) to test the effect of oLFS on spontaneously occurring seizures. During stimulation, we continuously recorded LFPs and videos. Recording sessions were divided into four sub-sessions: ‘pre’ - one hour before oLFS; ‘oLFS’ - one hour during oLFS; ‘post 1’ and ‘post 2’ - first hour and second hour after oLFS (Figure 1). For each frequency, we performed two trials on different days. For no-virus controls, only the ‘pre’ and ‘oLFS’ (1 Hz) sessions were done to check for light- or heat-induced effects.

Because in the intrahippocampal mouse model spontaneous recurrent seizures are frequent but mainly subclinical, whereas spontaneous behavioral seizures are rare, we additionally evoked these by 10 Hz photostimulation (25 ms pulse duration) (Figure 1) and assessed the effect of oLFS before or after the pro-convulsive 10 Hz stimulus. First, we systematically increased the stimulation duration (in 1-s steps) to determine the minimum duration sufficient to trigger a behavioral seizure for each animal (identification of seizure threshold). Each evoked seizure was manually inspected on the electrophysiological as well as behavioral level: we identified generic features as described by Jirsa et al. (2014) (Jirsa, Stacey, Quilichini, Ivanov, & Bernard, 2014) and motor symptoms according to the Racine scale (RS) (Racine, 1972). The identified seizure threshold was then validated for robust seizure induction for at least three times. In subsequent “preconditioning” oLFS sessions, photostimulation at 1 or 0.5 Hz was performed for 30 min before applying the pro-convulsive 10 Hz stimulus. For both frequencies, a minimum of three trials was performed on different days. In a subset of mice, ictogenic efficacy of the 10 Hz stimulus was tested again after the preconditioning experiments to exclude confounding effects of habituation.

### Perfusion and tissue preparation

Following the last recording session, mice were anesthetized (see above) 35 - 40 days after KA injection and transcardially perfused [0.9% saline followed by 4% paraformaldehyde in 0.1 M phosphate buffer (PB, pH 7.4)]. Following dissection, brains were post-fixated overnight, immersed in sucrose (25% in PB) overnight at 4°C for cryo-protection, shock-frozen in isopentane at −40°C and stored at −80°C. Brains were sectioned (coronal plane, 50 μm) with a cryostat (CM3050, Leica, Bensheim, GER). Slices were collected in 2x saline-sodium citrate buffer (2xSSC; 0.3 M NaCl, 0.03 M sodium citrate, pH 7.0).

### Fluorescent in situ hybridization

*Glutamic acid decarboxylase 67* (*Gad67*) mRNA was localized by FISH with digoxigenin (DIG)-labeled cRNA probes generated by *in vitro* transcription as described earlier (Kulik et al., 2003). Slices were hybridized with DIG-labeled antisense cRNA probes (see Supplementary material), and immunodetection of the DIG-labeled hybrids was performed with a peroxidase-conjugated anti-DIG antibody (1:2000; raised in sheep; Roche Diagnostics, Mannheim, GER). The fluorescence signal was developed with tyramide signal amplification (TSA) Plus Cyanine 3 System kit (PerkinElmer, Waltham, Massachusetts, USA) as described previously (Tulke et al., 2019).

### Immunohistochemistry

For immunofluorescence staining, free-floating sections were pre-treated in 10% normal horse serum (Vectorlabs, Burlingame, California, USA) in PB for one hour. Subsequently, slices were incubated first with guinea-pig anti-NeuN (1:500; Synaptic Systems, Göttingen, GER) overnight at 4 °C and then with a donkey anti-guinea-pig Cy5-conjugated antibody for 2.5 hours at RT (1:200, Jackson ImmunoResearch Laboratories Inc., West Grove, Pennsylvania, USA) followed by extensive rinsing in PB. Sections were mounted on glass slides with antifading mounting medium (DAKO, Hamburg, Germany).

### Image acquisition and analysis

Fluorescence composite images were taken with an AxioImager2 microscope using Plan-APOCHROMAT 5x or 10x objectives (Zeiss, Göttingen, GER). Exposure times (5x objective: Cy5-labeled NeuN, 500 ms; 10x objective: Cy3-labeled *Gad67* probe, 700 ms, Cy5-labeled NeuN, 5 s) were kept constant for each staining. The images were further processed with ZEN blue software (Zeiss).

To assess the extent of hippocampal sclerosis we quantified the volume of the dispersed granule cell layer (GCL) and cell loss in the hilus and CA1 along the septo-temporal axis of the hippocampus using Fiji ImageJ software (Schindelin et al., 2012). Here, masking was performed, since the evaluator was not aware of the respective kainate treatment. In detail, a region-of-interest (ROI) was drawn visually comprising the dispersed parts of the GCL using the ImageJ “polygon” function in each slice (around 50 per animal). Afterwards, the volume was calculated based on the area measured in each slice (values are given in mm^3^). For quantification of cell loss, the summed length of pyramidal cell-free gaps in the CA1 region was measured in each section using the “segmented line” function. Furthermore, hilar cell loss was quantified by automated detection (Cell Counter plugin) of NeuN^+^ cells (size parameter: 50-infinity pixel^2^) and *Gad67* mRNA^+^ interneurons (size parameter: 100-infinity pixel^2^) in the hilus of three dorsal sections for each animal. Here, we calculated the percentage of cell loss in the sclerotic hippocampus compared to the contralateral, non-sclerotic hippocampus (set to 100%). In detail, the Cy3 and Cy5 channels were split and individual images were converted to gray-scale. Images were background-subtracted by manual adjustment of the threshold and further processed using the “watershed” function, which separates overlapping cell bodies. A ROI was then defined for the hilus with the “polygon” function, and the respective cell density was calculated.

### Analysis of epileptiform activity

Recordings obtained from all electrodes were visually inspected for epileptiform activity. Animals that showed abnormal hippocampal atrophy were excluded from the analysis (n=1 for entorhinal fiber stimulation, n=3 for DGC stimulation). LFP data from the idHC site was then analyzed in detail using a custom algorithm (Heining et al., 2019). In the intrahippocampal KA mouse model, spontaneous recurrent seizures are evident as frequent bursts of high-amplitude sharp waves without behavioral symptoms (Riban et al., 2002). These bursts were classified according to their spike-load, hence, three categories of discharge patterns were identified (high-load, medium-load, and low-load bursts) as described by Heining et al. (2019) (Heining et al., 2019). To assess the severity of epileptiform activity within a recording, we calculated a ‘burst ratio’ as the quotient of the high-load burst duration and the total recording duration. The automatic detection of high-load bursts was verified by visual inspection. To assess epileptic burden of individual mice, the average burst ratio was calculated from a total of nine LFP recordings (15 hours) performed on different days (2x three hours before oLFS experiments, 6x one hour of ‘pre’ sessions of oLFS experiments and 1x three hours after oLFS experiments; Figure 1). In oLFS experiments, anti-epileptic effects were evaluated based on the burst ratio and the epileptic spike rate calculated for the respective sub-session (‘pre’, ‘oLFS’, ‘post 1’, post 2’). Individual trials in which a ‘pre’ recording had a burst ratio below 0.05 were excluded. Furthermore, we analyzed the magnitude of evoked responses during photostimulation using a 4^th^ order low pass Chebyshev filter with a cut-off frequency of 300 Hz to calculate the area under the curve (AUC, source code available). The time windows for the calculation of AUCs were set for each light pulse from −0.1 s to +0.2 s for oLFS (1, 0.5 and 0.2 Hz) and from −0.02 s to +0.06 s for the 10 Hz stimulation. To allow comparisons across animals, LFP data was z-scored and responses during high-load bursts were excluded from the calculation of the median AUC and polynomial fit. Recordings obtained from ivHC and cdHC sites were used to analyze the region-specific occurrence of epileptiform activity and to measure pulse latencies between recording sites during photostimulation sessions.

### Acute slice electrophysiology

In an additional set of experiments, mice were deeply anesthetized 21 to 28 days after KA (15 mM) and virus injection, perfused with 10 ml cold protective solution containing (in mM): 92 choline chloride, 30 NaHCO_3_, 2.5 KCl, 1.2 NaH_2_PO_4_, 25 D-glucose, 20 HEPES, 0.5 CaCl_2_, 5 Na-ascorbate acid, 2 Thiourea, 3 Na-Pyruvate, 10 MgCl_2_ and 12 acetylcysteine (oxygenated with 95% O_2_ / 5% CO_2_, 34°C) before dissection. Transverse acute hippocampal slices (300-350 µm) were obtained and incubated for 1 h at 34°C in a solution in which choline was replaced by 1 N-Methyl-D-glucamine (NMDG). Afterwards, slices were stored in artificial cerebrospinal fluid (ACSF, containing in mM: 125 NaCl, 25 NaHCO_3_, 2.5 KCl, 1.25 NaH_2_PO_4_, 25 D-glucose, 2 CaCl_2_ and 1 MgCl_2_ oxygenated with 95% O_2_ / 5% CO_2_) and supplemented with 12 mM N-acetylcysteine at RT. Whole-cell patch-clamp recordings were performed as previously described (Elgueta, Köhler, & Bartos, 2015) in the presence of GABA_A_ and GABA_B_ receptor blockers (10 µM gabazine and 2 µM CGP55845, respectively; 30-34°C; Multiclamp 700B amplifier (Molecular Devices, San José, California, USA); 5 kHz low-pass filter; sampling frequency 40 kHz). Stimulus generation, data acquisition and analysis were performed using custom-made programs written in Igor (WaveMetrics Inc., Portland, Oregon, USA).

Recording pipettes were filled with a solution containing (in mM): 140 K-Gluconate, 4 KCl, 10 HEPES, 2 MgCl_2_, 2 Na_2_ATP, 10 EGTA, 0.125 Alexa-Fluor 488 and 0.15 % biocytin (pH = 7.2; 290-310 mOsm), that resulted in pipette resistances of 4-6 MΩ. Series resistances between (8 – 20 MΩ) were compensated using bridge balance in current-clamp and were left uncompensated in voltage-clamp. For loose-patch experiments and extracellular stimulations, pipettes were filled with a HEPES-buffered ACSF (containing in mM: ACSF (see above), 135 NaCl, 5.4 KCl, 1.8 CaCl_2_, 1 MgCl_2_, 5 HEPES). Extracellular stimulation was performed using a stimulus isolator (Isopulser) with pipettes (∼1 MΩ) placed in the middle molecular layer where the virus was expressed. Five pulses (50 Hz, 0.1–0.3 ms, 20–100 V) were evoked and a minimum of ten trials was used to calculate the overall discharge probability. The rheobase was measured with 1 s-long current injections increasing with 20 pA steps. Series resistance, cell capacitance (C_m_) and membrane resistance (R_m_) were calculated from −10 mV pulses. 10 min photostimulation was performed using full-field blue light pulses (473 nm; 50 ms pulse duration; LED p2000, CoolLED, Andover, UK).

### Statistical analysis

Data were tested for significant differences with Prism 7 software (GraphPad Software Inc.). Comparisons of two groups were performed with a paired (comparisons within animals) or unpaired (comparisons between animals) Student’s t-test. Multiple-group comparisons were calculated with an ordinary or repeated measure (RM) one-way ANOVA followed by a Tukey’s *post hoc* test. Significance thresholds were set to: \**p*<0.05, \*\**p*<0.01 and \*\*\**p*<0.001. For all values, mean and standard error of the mean (SEM) are given, unless otherwise reported. Correlations were tested using Pearson’s correlation (slope significantly non-zero, confidence interval (CI) 95 %).

## Data availability statement

The LFP dataset has been made available https://osf.io/uk94m/. The source code and user information for the seizure detection algorithm can be obtained from Ulrich Egert upon request. Contact

## Acknowledgements

We thank Piret Kleis and Lea Hüper for support with data analysis and Andrea Djie-Maletz for excellent technical assistance. We thank Dr. Antje Kilias for critical discussions and valuable input.

## Author contributions

EP, Data curation, Formal analysis, Investigation, Visualization, Methodology, Writing— original draft preparation, Writing—review and editing; CE, Data curation, Formal analysis, Investigation, Visualization, Methodology, Writing—original draft preparation, Writing—review and editing; KH, Formal analysis, Software, Investigation, Visualization, Writing—review and editing; DV, Formal analysis, Investigation, Visualization, Writing— review and editing; CO, Data curation; UH, Validation, Methodology, Writing—review and editing; MB, Funding acquisition; UE, Funding acquisition, Methodology, Writing—review and editing; PJ, Conceptualization, Supervision, Validation, Data curation, Formal analysis, Investigation, Methodology, Writing—review and editing; CAH, Conceptualization, Supervision, Funding acquisition, Writing—review and editing

## Supplementary Materials and Methods

### Fluorescent in situ hybridization

Brain slices were pre-treated in a 1:1 mixture of hybridization buffer [50% formamide, 4xSSC, 5% dextransulfate, 250 μg/ml heat-denatured salmon sperm DNA, 200 µl yeast t-RNA, 1% Denhardt’s-reagent (Sigma-Aldrich, Steinheim, GER)] and 2xSSC at RT for 15 minutes. Subsequently, the slices were pre-hybridized in hybridization buffer for 60 min at 45°C, followed by addition of DIG-labeled antisense or sense *Gad67* cRNA probe (100 ng/ml) and incubated overnight at 45°C. Slices were washed in 2xSSC for 2 x 15 min at RT and then successively rinsed at 55°C for 15 min in 2x SSC with 50% formamide; 0.1x SSC with 50% formamide and twice in 0.1x SSC alone. Then the slices were rinsed in 0.1 M Tris-buffered saline (TBS) for 2x 10 min and transferred to the blocking buffer [1% blocking reagent (Roche Diagnostics, Mannheim, GER) in TBS] for 60 min at RT. Slices were pre-hybridized at 45°C, followed by addition of DIG-labeled antisense or sense *Gad67* cRNA probe (100 ng/ml) and incubated overnight at 45°C. For fluorescent detection, tissue sections were treated with a horseradish peroxidase-conjugated DIG antibody (1:2000, raised in sheep; Roche Diagnostics) overnight at 4°C and developed in the presence of amplification buffer and tyramide working solution (1:50) for 6 min in the dark, using the Tyramide Signal Amplification (TSA) Plus Cyanine 3 System kit (PerkinElmer, Waltham, USA)). The staining-reaction was stopped by rinsing in TBS for 3x 5 min and 1x 15 min. Slices were kept in the dark for further immunofluorescence staining.

**Supplementary Figure 1.**
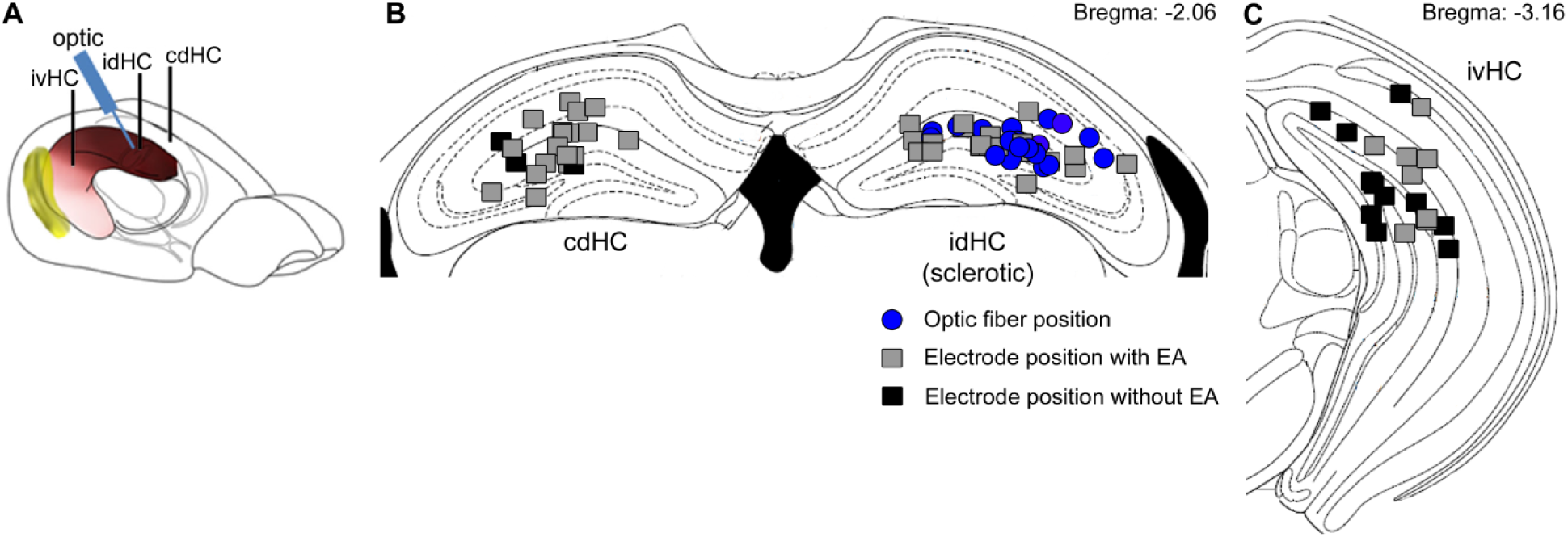
Positions of implanted electrodes and optic fibers for all animals included in the study. **(A)** Implantation scheme. Electrode positions for all animals (n=21) were histologically validated and were located at three different recording sites: cdHC, contralateral dorsal hippocampus; idHC, ipsilateral dorsal hippocampus and ivHC, ipsilateral ventral hippocampus. The optic fiber was placed in a 30° angle at the idHC site. **(B)** Positions of the optic fiber (blue circle) and electrodes in the dorsal region, color-coded for the detection of epileptiform activity (EA, grey square) or no EA (black square) on the respective electrode. **(C)** Ipsilateral ventral electrode positions.

**Supplementary Figure 2.**
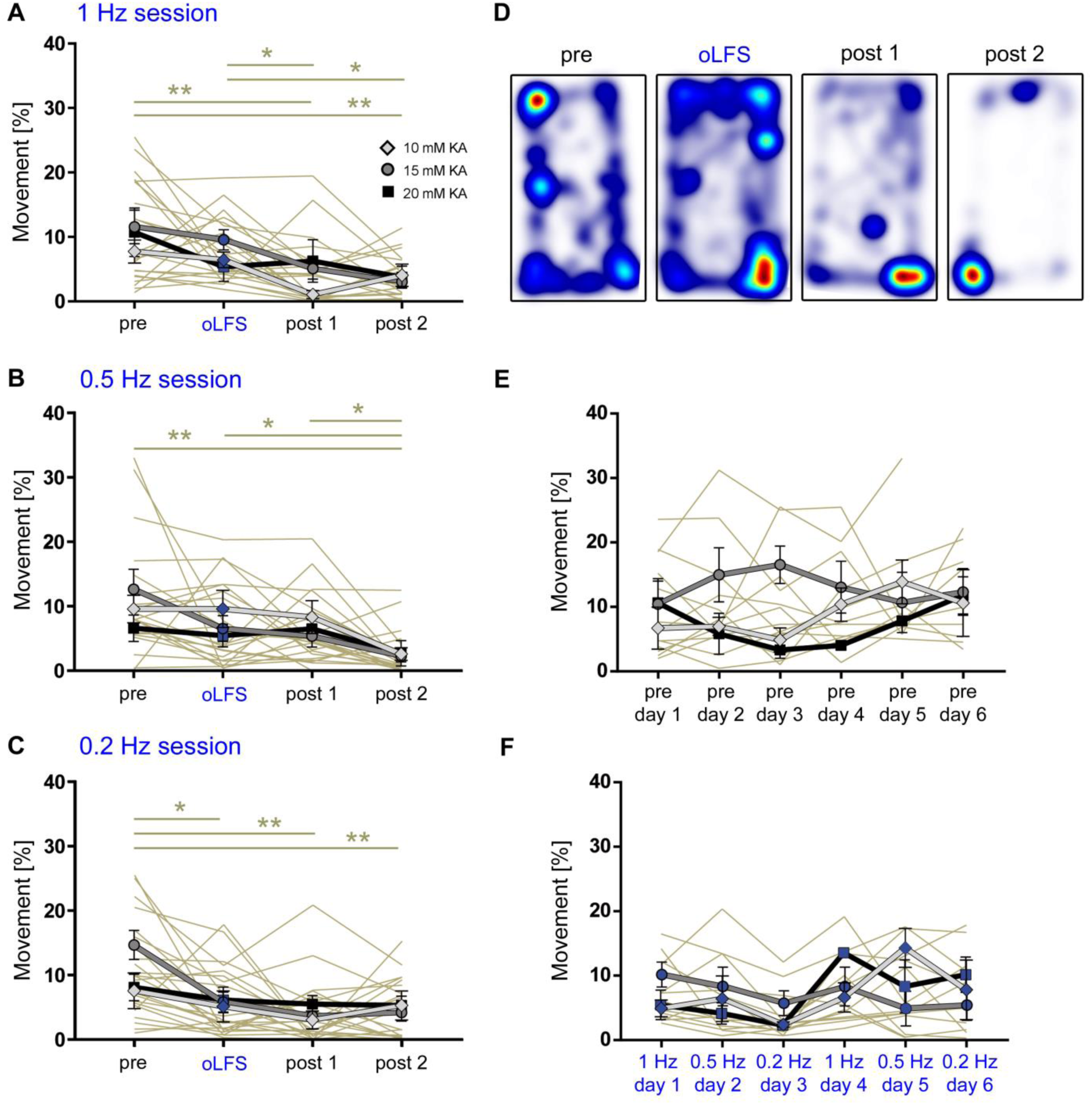
Movement analysis of chronically epileptic mice during LFP recordings. Freely-moving animals were video tracked, running phases were automatically detected and quantified as time spent running (>4 cm/s). **(A – C)** Analysis of all sub-session clearly reveals that all mice, independent of the KA concentration, exhibit normal running behavior that declines gradually during the recording time of four hours. **(D)** Representative heat map of a 1 Hz session with the corresponding ‘pre’ and ‘post’ recordings. Warm colors indicate places of longer stays during each sub-session. **(E, F)** Mice don’t change their running behavior during oLFS experiments (six days in a row) in the ‘pre’ or the ‘oLFS’ sessions. Hence there is no adaptation to the environment across days and no difference between stimulation frequencies. Individual values are presented in Supplementary Table S3. Single sessions (beige) are used to calculate the RM one-way ANOVA; Tukey’s multiple comparison test (all KA concentrations merged); *p<0.05 and **p<0.01. Values are given as mean ± SEM.

**Supplementary Figure 3.**
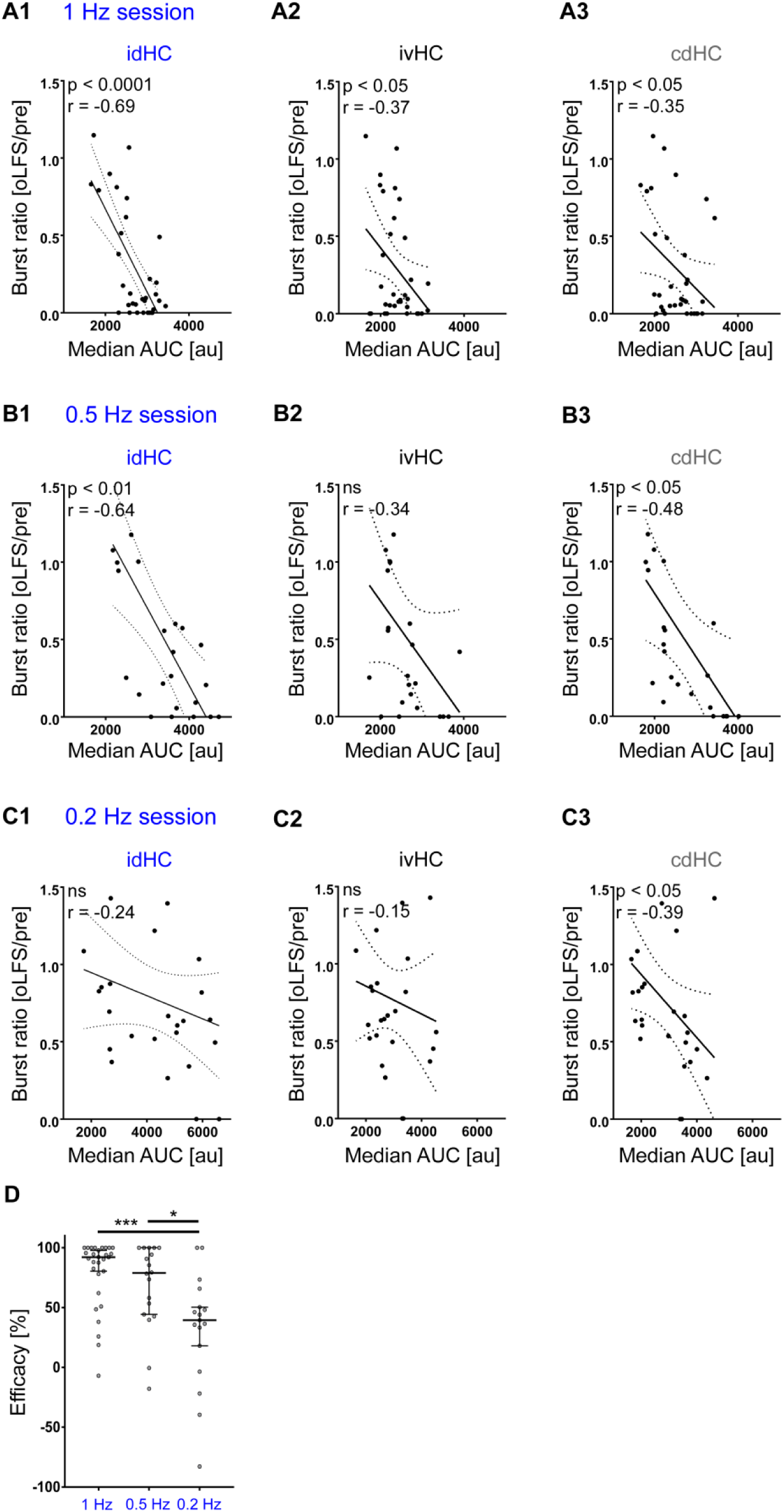
Stimulation efficacy of oLFS depends on the magnitude of evoked responses. Linear regression analysis of the stimulation efficacy (oLFS/pre burst ratio) and the median AUC of the respective session at all electrode positions. **(A1-3)** 1 Hz sessions. **(B1-3)** 0.5 Hz sessions. **(C1-3)** 0.2 Hz sessions. **(D)** For comparison of the stimulation efficacy between frequencies, we consider only sessions with a cellular response (at idHC) within the range of the standard deviation. 1 Hz oLFS is the most effective stimulation paradigm, but 0.5 and 0.2 Hz also elicit remarkable anti-ictogenic effects. One-way ANOVA; Dunn’s multiple comparison test; *p<0.05, ***p<0.001. Values are given as median ± 95% CI.

**Supplementary Figure 4.**
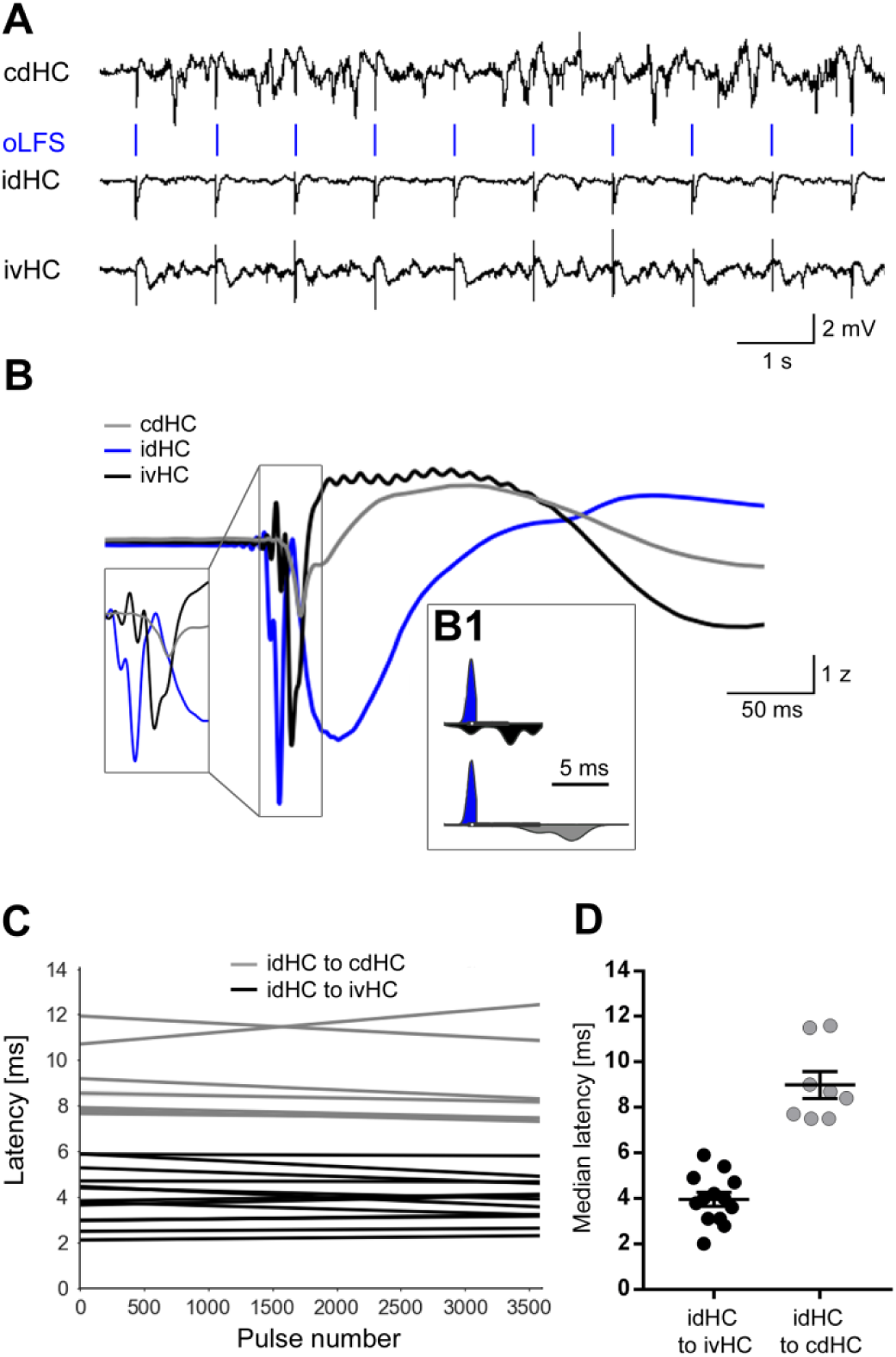
Local oLFS leads to delayed responses in other regions of both hippocampi. **(A)** Representative LFP traces of all three electrodes during 1 Hz oLFS. Local stimulation of DGCs via entorhinal afferents in the idHC evokes population spikes also in the ivHC and cdHC. **(B)** Population spikes occur first in the idHC (blue), followed by ivHC (black) and cdHC (grey) (representative example). **(B1)** Distribution of spike times for 3600 responses (one hour, 1 Hz stimulation) at the three electrode positions. **(C)** Robust linear regression shows stable latencies from idHC to ivHC (black, n=12 sessions from 8 animals) and cdHC regions (grey, n=8 sessions from 7 animals) over one hour oLFS. **(D)** Population spikes occur with a median latency of 4 ms and 8.5 ms in the ivHC and cdHC region, respectively.

**Supplementary Figure 5.**
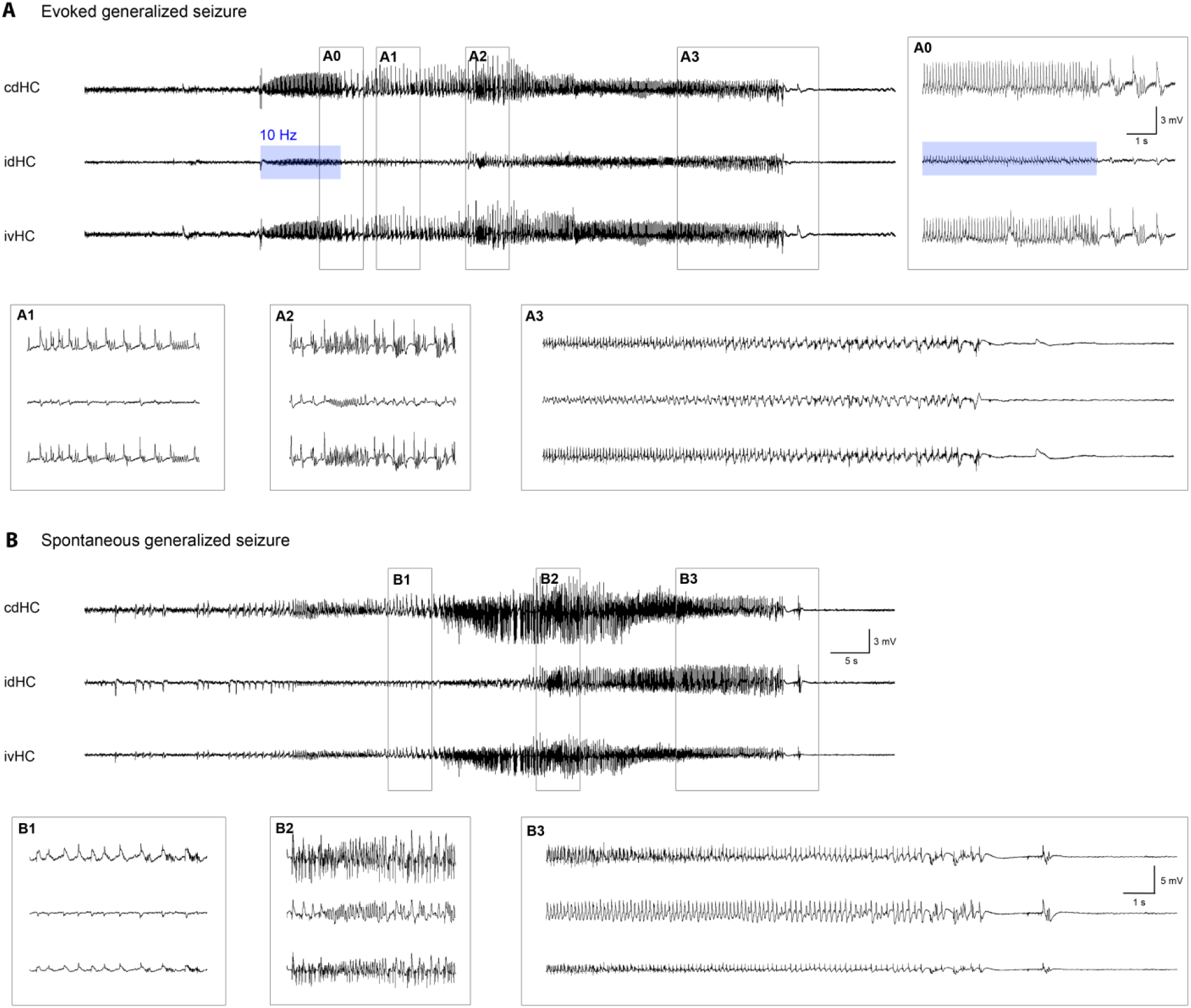
Comparison of spontaneous and evoked generalized seizures. **(A, B)** Representative LFP traces of an optogenetically evoked and a spontaneous generalized seizure from the same animal. **(A0)** High-amplitude epileptic spikes emerge in addition to the evoked potentials during stimulation and rhythmic activity persists after 10 Hz stimulation has been stopped. Both seizures consist of the same building blocks (**(A1, B1)** spike-and-wave events; **(A2, B2)** fast discharges; **(A3, B3)** increasing inter-spike-intervals and subsequent termination) with similar dynamics. This consistency is evident for all mice (n=5) in which both spontaneous and evoked generalized seizures could be detected.

## References

Abrahamsson, T., Gustafsson, B., & Hanse, E. (2005). Synaptic fatigue at the naive perforant path-dentate granule cell synapse in the rat. J Physiol, 569(3), 737–750. https://doi.org/10.1113/jphysiol.2005.097725

Avoli, M., de Curtis, M., & Köhling, R. (2013). Does interictal synchronization influence ictogenesis? Neuropharmacology, 69, 37–44. https://doi.org/10.1016/j.neuropharm.2012.06.044.Does

Blümcke, I., Zuschratter, W., Schewe, J. C., Suter, B., Lie, A. A., Riederer, B. M., … Wiestler, O. D. (1999). Cellular pathology of hilar neurons in Ammon’s horn sclerosis. J Comp Neurol, 414(4), 437–453. https://doi.org/10.1002/(SICI)1096-9861(19991129)414:4<437::AID-CNE2>3.0.CO;2-3

Boëx, C., Seeck, M., Vulliémoz, S., Rossetti, A. O., Staedler, C., Spinelli, L., … Pollo, C. (2011). Chronic deep brain stimulation in mesial temporal lobe epilepsy. Seizure, 20(6), 485–490. https://doi.org/10.1016/j.seizure.2011.03.001

Bouilleret, V., Ridoux, V., Depaulis, A., Marescaux, C., Nehlig, A., & Le Gal La Salle, G. (1999). Recurrent seizures and hippocampal sclerosis following intrahippocampal kainate injection in adult mice: Electroencephalography, histopathology and synaptic reorganization similar to mesial temporal lobe epilepsy. Neuroscience, 89(3), 717–729. https://doi.org/10.1016/S0306-4522(98)00401-1

Chang, W. C., Kudlacek, J., Hlinka, J., Chvojka, J., Hadrava, M., Kumpost, V., … Jiruska, P. (2018). Loss of neuronal network resilience precedes seizures and determines the ictogenic nature of interictal synaptic perturbations. Nat Neurosci, 21(12), 1742–1752. https://doi.org/10.1038/s41593-018-0278-y

De Curtis, M., & Avanzini, G. (2001). Interictal spikes in focal epileptogenesis. Prog Neurobiol, 63(5), 541–567. https://doi.org/10.1016/S0301-0082(00)00026-5

De Curtis, M., Librizzi, L., & Biella, G. (2001). Discharge threshold is enhanced for several seconds after a single interictal spike in a model of focal epileptogenesis. Eur J Neurosci, 14(1), 174–178. https://doi.org/10.1046/j.0953-816X.2001.01637.x

Elgueta, C., Köhler, J., & Bartos, M. (2015). Persistent discharges in dentate gyrus perisoma-inhibiting interneurons require hyperpolarization-activated cyclic nucleotide-gated channel activation. J Neurosci, 35(10), 4131–4139. https://doi.org/10.1523/JNEUROSCI.3671-14.2015

Engel, J. (2001). Mesial temporal lobe epilepsy: What have we learned? The Neuroscientist, 7(4), 340–352. https://doi.org/10.1177/107385840100700410

Feng, G., Mellor, R. H., Bernstein, M., Keller-Peck, C., Nguyen, Q. T., Wallace, M., … Sanes, J. R. (2000). Imaging neuronal subsets in transgenic mice expressing multiple spectral variants of GFP. Neuron, 28(1), 41–51. https://doi.org/10.1016/S0896-6273(00)00084-2

Fisher, R. S., & Velasco, A. L. (2014). Electrical brain stimulation for epilepsy. Nat Rev Neurol, 10(5), 261–270. https://doi.org/10.1038/nrneurol.2014.59

Gonzalez, J., Morales, I. S., Villarreal, D. M., & Derrick, B. E. (2014). Low-frequency stimulation induces long-term depression and slow onset long-term potentiation at perforant path-dentate gyrus synapses in vivo. J Neurophys, 111(6), 1259–1273. https://doi.org/10.1152/jn.00941.2012

Häussler, U., Bielefeld, L., Froriep, U. P., Wolfart, J., & Haas, C. A. (2012). Septotemporal position in the hippocampal formation determines epileptic and neurogenic activity in temporal lobe epilepsy. Cereb Cortex, 22(1), 26–36. https://doi.org/10.1093/cercor/bhr054

Häussler, U., Rinas, K., Kilias, A., Egert, U., & Haas, C. A. (2016). Mossy fiber sprouting and pyramidal cell dispersion in the hippocampal CA2 region in a mouse model of temporal lobe epilepsy. Hippocampus, 26(5), 577–588. https://doi.org/10.1002/hipo.22543

Heining, K., Kilias, A., Janz, P., Häussler, U., Kumar, A., Haas, C. A., & Egert, U. (2019). Bursts with high and low load of epileptiform spikes show context-dependent correlations in epileptic mice. ENeuro, 6(5), 1–14. https://doi.org/10.1523/ENEURO.0299-18.2019

Heinrich, C., Nitta, N., Flubacher, A., Mueller, M., Fahrner, A., Kirsch, M., … Haas, C. A. (2006). Reelin deficiency and displacement of mature neurons, but not neurogenesis, underlie the formation of granule cell dispersion in the epileptic hippocampus. J Neurosci, 26(17), 4701–4713. https://doi.org/10.1523/JNEUROSCI.5516-05.2006

Janz, P., Hauser, P., Heining, K., Nestel, S., Kirsch, M., Egert, U., & Haas, C. A. (2018). Position- and time-dependent Arc expression links neuronal activity to synaptic plasticity during epileptogenesis. Front Cell Neurosci, 12(8), 1–18. https://doi.org/10.3389/fncel.2018.00244

Janz, P., Savanthrapadian, S., Häussler, U., Kilias, A., Nestel, S., Kretz, O., … Haas, C. A. (2017). Synaptic remodeling of entorhinal input contributes to an aberrant hippocampal network in temporal lobe epilepsy. Cereb Cortex, 27(3), 2348–2364. https://doi.org/10.1093/cercor/bhw093

Janz, P., Schwaderlapp, N., Heining, K., Häussler, U., Korvink, J. G., von Elverfeldt, D., … Haas, C. A. (2017). Early tissue damage and microstructural reorganization predict disease severity in experimental epilepsy. ELife, 6, e25742. https://doi.org/10.7554/eLife.25742

Jirsa, V. K., Stacey, W. C., Quilichini, P. P., Ivanov, A. I., & Bernard, C. (2014). On the nature of seizure dynamics. Brain, 137(8), 2210–2230. https://doi.org/10.1093/brain/awu133

Kile, K. B., Tian, N., & Durand, D. M. (2010). Low frequency stimulation decreases seizure activity in a mutation model of epilepsy. Epilepsia, 51(9), 1745–1753. https://doi.org/10.1111/j.1528-1167.2010.02679.x.Low

Kokaia, M., Andersson, M., & Ledri, M. (2013). An optogenetic approach in epilepsy. Neuropharmacology, 69, 89–95. https://doi.org/10.1016/j.neuropharm.2012.05.049

Krook-Magnuson, E., Armstrong, C., Bui, A., Lew, S., Oijala, M., & Soltesz, I. (2015). In vivo evaluation of the dentate gate theory in epilepsy. J Physiol, 593(10), 2379–88. https://doi.org/10.1113/JP270056

Krook-Magnuson, E., Armstrong, C., Oijala, M., & Soltesz, I. (2013). On-demand optogenetic control of spontaneous seizures in temporal lobe epilepsy. Nat Commun, 4, 1376. https://doi.org/10.1038/ncomms2376

Krook-Magnuson, E., Szabo, G. G., Armstrong, C., Oijala, M., & Soltesz, I. (2014). Cerebellar directed optogenetic intervention inhibits spontaneous hippocampal seizures in a mouse model of temporal lobe epilepsy. ENeuro, 1(1), 1–15. https://doi.org/10.1523/ENEURO.0005-14.2014

Kulik, Á., Vida, I., Luján, R., Haas, C. A., López-Bendito, G., Shigemoto, R., & Frotscher, M. (2003). Subcellular localization of metabotropic GABA(B) receptor subunits GABA(B1a/b) and GABA(B2) in the rat hippocampus. J Neurosci, 23(35), 11026– 11035. https://doi.org/10.1523/jneurosci.23-35-11026.2003

Ladas, T. P., Chiang, C.-C., Gonzalez-Reyes, L. E., Nowak, T., & Durand, D. M. (2015). Seizure reduction through interneuron-mediated entrainment using low frequency optical stimulation. Exp Neurol, 2(74), 120–132. https://doi.org/10.1126/scisignal.274pe36.Insulin

Ledri, M., Madsen, M. G., Nikitidou, L., Kirik, D., & Kokaia, M. (2014). Global optogenetic activation of inhibitory interneurons during epileptiform activity. J Neurosci, 34(9), 3364–3377. https://doi.org/10.1523/JNEUROSCI.2734-13.2014

Lévesque, M., Chen, L., Etter, G., Shiri, Z., Wang, S., Williams, S., & Avoli, M. (2019). Paradoxical effects of optogenetic stimulation in mesial temporal lobe epilepsy. Ann Neurol, 86(5), 714–728. https://doi.org/10.1002/ana.25572

Li, M. C. H., & Cook, M. J. (2018). Deep brain stimulation for drug-resistant epilepsy. Epilepsia, 59(2), 273–290. https://doi.org/10.1111/epi.13964

Lim, S.-N., Lee, C.-Y., Lee, S.-T., Tu, P.-H., Chang, B.-L., Lee, C.-H., … Wu, T. (2016). Low and high frequency hippocampal stimulation for drug-resistant mesial temporal lobe epilepsy. Neuromodulation, 19(4), 365–372. https://doi.org/10.1111/ner.12435

Lu, Y., Zhong, C., Wang, L., Wei, P., He, W., Huang, K., … Wang, L. (2016). Optogenetic dissection of ictal propagation in the hippocampal-entorhinal cortex structures. Nat Commun, 7, 10962. https://doi.org/10.1038/ncomms10962

Marx, M., Haas, C. A., & Häussler, U. (2013). Differential vulnerability of interneurons in the epileptic hippocampus. Front Cell Neurosci, 7, 167. https://doi.org/10.3389/fncel.2013.00167

Meyer, M., Kienzler-Norwood, F., Bauer, S., Rosenow, F., & Norwood, B. A. (2016). Removing entorhinal cortex input to the dentate gyrus does not impede low frequency oscillations, an EEG-biomarker of hippocampal epileptogenesis. Sci Rep, 6, 2–10. https://doi.org/10.1038/srep25660

Mohan, M., Keller, S., Nicolson, A., Biswas, S., Smith, D., Farah, J. O., … Wieshmann, U. (2018). The long-term outcomes of epilepsy surgery. PLoS ONE, 13(5), 1–16. https://doi.org/10.1371/journal.pone.0196274

Muldoon, S. F., Villette, V., Tressard, T., Malvache, A., Reichinnek, S., Bartolomei, F., & Cossart, R. (2015). GABAergic inhibition shapes interictal dynamics in awake epileptic mice. Brain, 138(10), 2875–2890. https://doi.org/10.1093/brain/awv227

Osawa, S., Iwasaki, M., Hosaka, R., Matsuzaka, Y., & Tomita, H. (2013). Optogenetically induced seizure and the longitudinal hippocampal network dynamics. PLoS ONE, 8(4), 1–14. https://doi.org/10.1371/journal.pone.0060928

Pallud, J., Häussler, U., Hamelin, S., Devaux, B., Deransart, C., & Depaulis, A. (2011). Dentate gyrus and hilus transection blocks seizure propagation and granule cell dispersion in a mouse model for mesial temporal lobe epilepsy. Hippocampus, 21(3), 334–343. https://doi.org/10.1002/hipo.20795

Racine, R. J. (1972). Modification of seizure activity by electrical stimulation. II. Motor seizure. Electroencephalogr Clin Neurophysiol., 32(3), 281–94.

Rashid, S., Pho, G., Czigler, M., Werz, M. A., & Durand, D. M. (2012). Low frequency stimulation of hippocampal commissures reduces seizures in chronic rat model of temporal lobe epilepsy. Epilepsia, 53(1), 147–156. https://doi.org/10.1111/j.1528-1167.2011.03348.x.Low

Riban, V., Bouilleret, V., Pham-Le, B. T., Fritschy, J. M., Marescaux, C., & Depaulis, A. (2002). Evolution of hippocampal epileptic activity during the development of hippocampal sclerosis in a mouse model of temporal lobe epilepsy. Neuroscience, 112(1), 101–111. https://doi.org/10.1016/S0306-4522(02)00064-7

Ryvlin, P., & Kahane, P. (2005). The hidden causes of surgery-resistant temporal lobe epilepsy: extratemporal or temporal plus? Curr Opin Neurol, 18(2), 125–127. https://doi.org/10.1097/01.wco.0000162852.22026.6f

Schindelin, J., Arganda-Carreras, I., Frise, E., Kaynig, V., Pietzsch, T., Preibisch, S., … Cardona, A. (2012). Fiji - an open source platform for biological image analysis. Nat Methods, 9(7), 676–682. https://doi.org/10.1038/nmeth.2019.Fiji

Shiri, Z., Lévesque, M., Etter, G., Manseau, F., Williams, S., & Avoli, M. (2017). Optogenetic low-frequency stimulation of specific neuronal populations abates ictogenesis. Neurobiol Dis, 37(11), 2999–3008. https://doi.org/10.1523/JNEUROSCI.2244-16.2017

Thom, M. (2014). Review: Hippocampal sclerosis in epilepsy: A neuropathology review. Neuropathol Appl Neurobiol, 40(5), 520–543. https://doi.org/10.1111/nan.12150

Tulke, S., Haas, C. A., & Häussler, U. (2019). Expression of brain-derived neurotrophic factor and structural plasticity in the dentate gyrus and CA2 region correlate with epileptiform activity. Epilepsia, 60(6), 1234–1247. https://doi.org/10.1111/epi.15540

Velasco, A. L., Velasco, F., Velasco, M., Trejo, D., Castro, G., & Carrillo-Ruiz, J. D. (2007). Electrical stimulation of the hippocampal epileptic foci for seizure control: A double-blind, long-term follow-up study. Epilepsia, 48(10), 1895–1903. https://doi.org/10.1111/j.1528-1167.2007.01181.x

Xu, Z., Wang, Y., Chen, B., Xu, C., Wu, X., Wang, Y., … Chen, Z. (2016). Entorhinal principal neurons mediate brain-stimulation treatments for epilepsy. EBioMedicine, 14, 148–160. https://doi.org/10.1016/j.ebiom.2016.11.027

Young, C. C., Stegen, M., Bernard, R., Müller, M., Bischofberger, J., Veh, R. W., … Wolfart, J. (2009). Upregulation of inward rectifier K+ (Kir2) channels in dentate gyrus granule cells in temporal lobe epilepsy. J Physiol, 587(17), 4213–4233. https://doi.org/10.1113/jphysiol.2009.170746

Zeineh, M. M., Palomero-Gallagher, N., Axer, M., Gräßel, D., Goubran, M., Wree, A., … Zilles, K. (2017). Direct visualization and mapping of the spatial course of fiber tracts at microscopic resolution in the human hippocampus. Cereb Cortex, 27(3), 1779–1794. https://doi.org/10.1093/cercor/bhw010

Zhao, M., Alleva, R., Ma, H., Daniel, A. G. S., & Schwartz, T. H. (2015). Optogenetic tools for modulating and probing the epileptic network. Epilepsy Res., 116, 15–26. https://doi.org/10.1016/j.eplepsyres.2015.06.010

